# A combinatorial neural code for long-term motor memory

**DOI:** 10.1101/2024.06.05.597627

**Authors:** Jae-Hyun Kim, Kayvon Daie, Nuo Li

## Abstract

Motor skill repertoire can be stably retained over long periods, but the neural mechanism underlying stable memory storage remains poorly understood. Moreover, it is unknown how existing motor memories are maintained as new motor skills are continuously acquired. Here we tracked neural representation of learned actions throughout a significant portion of a mouse’s lifespan, and we show that learned actions are stably retained in motor memory in combination with context, which protects existing memories from erasure during new motor learning. We used automated home-cage training to establish a continual learning paradigm in which mice learned to perform directional licking in different task contexts. We combined this paradigm with chronic two-photon imaging of motor cortex activity for up to 6 months. Within the same task context, activity driving directional licking was stable over time with little representational drift. When learning new task contexts, new preparatory activity emerged to drive the same licking actions. Learning created parallel new motor memories while retaining the previous memories. Re-learning to make the same actions in the previous task context re-activated the previous preparatory activity, even months later. At the same time, continual learning of new task contexts kept creating new preparatory activity patterns. Context-specific memories, as we observed in the motor system, may provide a solution for stable memory storage throughout continual learning. Learning in new contexts produces parallel new representations instead of modifying existing representations, thus protecting existing motor repertoire from erasure.

## Introduction

In our lifetime we stably retain a myriad of motor skills. How are learned actions stored in motor memory? In sensory systems, neurons learn to represent conjunctions of features extracted by sensory organs ^1–4^. In contrast to the representational view of the sensory systems, the prevailing view of the motor system is the dynamical systems framework ^5,6^, in which neural activity is not a function of behavioral variables but internally generated activity state. In the motor cortex, specific learned actions are evoked by distinct patterns of preparatory activity ^7–11^ (Fig. 1a), which reflect persistent activity states supported by recurrent networks in motor cortex and connected brain areas ^6,12–15^. Preparatory activity is thought to provide the initial conditions for the ensuing dynamics dictating movement execution ^6,16–20^, but its relationship to subsequent action remains obscure ^21–23^. For example, are preparatory activity states linked to subsequent movement execution and therefore fixed for actions with identical kinematics? Alternatively, might preparatory activity encode other cognitive variables associated with learned actions beyond the movement itself ^10,11,24–26^?

**Figure 1.**
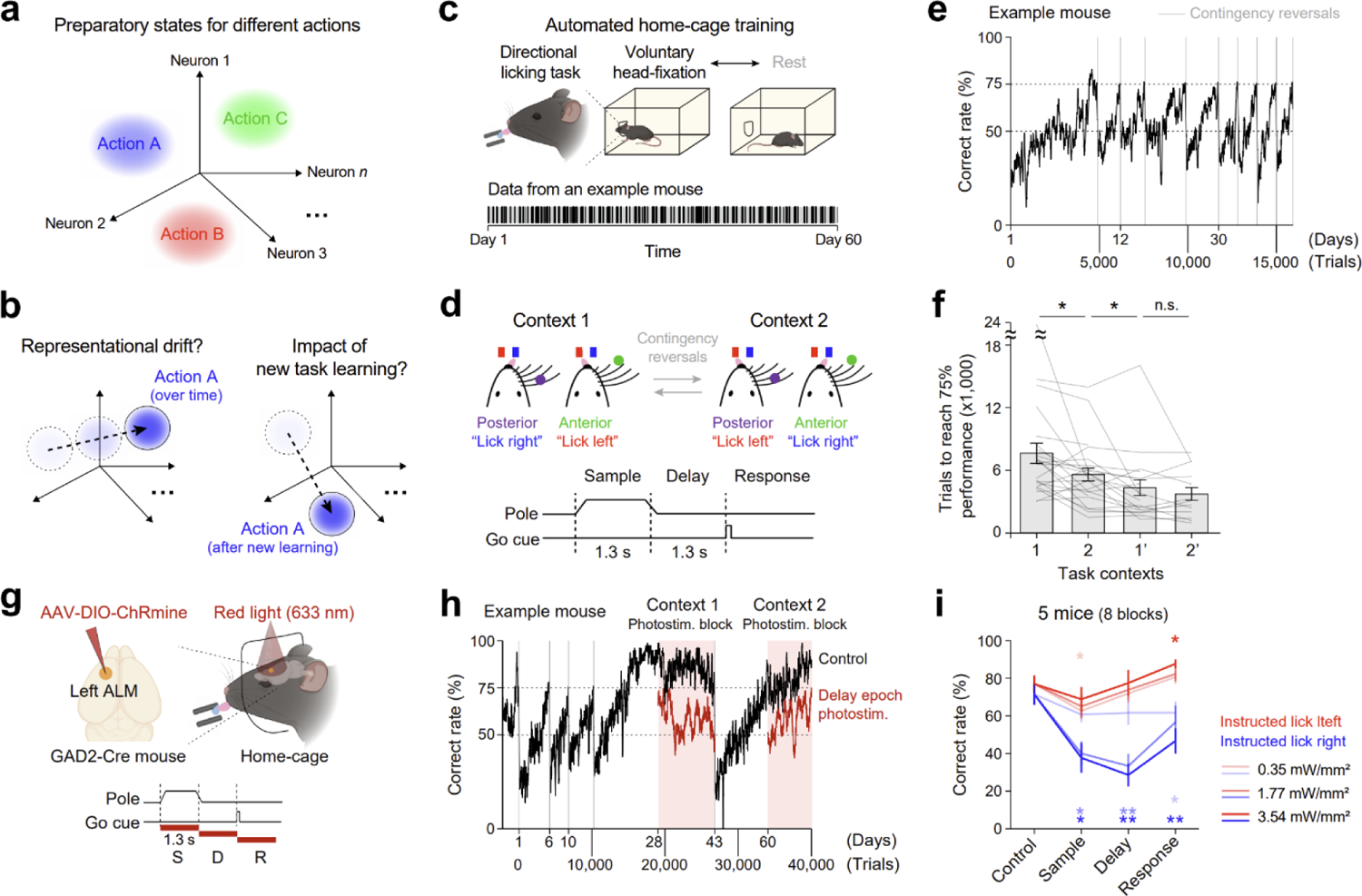
A behavior paradigm for continual learning that requires ALM. **a.** Schematic of preparatory states for different actions in activity space. **b.** Possible outcomes of preparatory states for the same action over time and across new task learning involving the action. **c.** Automated home-cage training. Top, schematic of mice living in the home-cage system voluntarily engage in head-fixation and learn directional lick tasks. Bottom, chronological behavioral data from an example mouse. Dark bands, epochs of voluntary head-fixation; gray, rest. **d.** Directional licking tasks. Mice discriminate pole position during the sample epoch and report position using lick left or lick right after the delay epoch. Sensorimotor contingency is reversed across task contexts. **e.** Behavior performance from an example mouse learning multiple rounds of contingency reversals. Gray vertical lines indicate the onset of contingency reversals, introduced when performance is above 75%. Averaging window, 100 trials. **f.** The number of trials to reach 75% correct performance during initial learning and subsequent reversal learnings (mean ± s.e.m.). Individual lines show individual mice. Mice used for in-cage optogenetic (5 mice), imaging (13 mice), and behavior testing only (5 mice) are combined. Learning task context 1 vs. 2, *P=0.0286 (23 mice); learning task context 2 vs. 1’, *P=0.0431 (19 mice); learning task context 1’ vs. 2’, P=0.3425 (13 mice). Paired t-test. **g.** Top, optogenetic approach to silence ALM activity during task performance in home-cage. Bottom, task and photoinhibition timelines. Photostimulation during the sample (S), delay (D), or response epoch (R). Power, 0.35, 1.77, 3.54 mW/mm^2^. **h.** Behavior performance from an example mouse during ALM photoinhibition after multiple rounds of contingency reversals. Black, control trials. Red, photoinhibition during the delay epoch (3.54 mW/mm^2^). Light red shade, photoinhibition blocks. **i.** Behavior performance during ALM photoinhibition (mean ± s.e.m.). Trial types by instructed lick direction. Left ALM photostimuluation. Sample epoch, instructed lick right, *P=0.0248, F=0.7574 (1.77 mW/mm^2^), *P=0.0349, F=0.8402 (3.54 mW/mm^2^); instructed lick left, *P=0.0360, F=1.0334 (0.35 mW/mm^2^). Delay epoch, instructed lick right, **P=0.0054, F=0.7212 (1.77 mW/mm^2^), **P=0.0012, F=0.3909 (3.54 mW/mm^2^). Response epoch, instructed lick right, *P=0.0249, F=0.4940 (0.35 mW/mm^2^), **P=0.0093, F=0.6863 (3.54 mW/mm^2^); instructed lick left, *P=0.0423, F=1.0702 (3.54 mW/mm^2^). P values by two-tailed t-test against control.

A related question is how learned actions are maintained by motor circuits over time. Motor cortex circuits exhibit considerable plasticity ^27–35^, and motor cortex activity undergoes systematic reconfigurations during motor learning ^9,11,31,36–49^. Given this plasticity, the neural mechanism underlying motor memory storage is unclear. Recent studies propose memory storage mechanisms based on unstable representations ^50^: in a redundant neural network in which multiple network configurations produce the same output, activity patterns leading to the same motor output can change over time ^51,52^, allowing the network to explore multiple solutions to solve the same task ^50,53^. For example, if a pattern of activity drives our speech of the word “cat”, would it be the same pattern of activity when we utter the word “cat” a year later (Fig. 1b, left)? This question remains under-explored as neural encoding of movement has mostly been examined across hours or days ^11,31,36,40,41,54,55^, and rarely beyond one month ^56–58^.

Moreover, it is unknown how existing motor memories are protected from modifications by continual learning of new motor skills, as new experience can overwrite previously learned representations. Theories of learning posit a modular approach, where multiple parallel motor memories are formed for distinct contexts ^24,25,59,60^, thus previously learned motor skills are retained while new learning takes place in separate modules. Yet, neurophysiological studies of motor learning have mostly examined single tasks. It remains poorly understood how neural representation of an action is formed and maintained when we learn to utilize the same action in different contexts, e.g., learning to speak the word “cat” in different sentences (Fig. 1b, right).

To address these questions, we used automated home-cage training to establish a continual learning paradigm in which mice learned to perform directional licking in different task contexts. In-cage optogenetics shows that learned directional licking is dependent on preparatory activity in anterior lateral motor cortex (ALM) ^19,61,62^. We used two-photon calcium imaging to chronically track ALM activity across continual learning for multiple months. Within the same task context, the preparatory activity encoding directional licking exhibited little representational drift over time. As mice learned new task contexts, new preparatory activity emerged to drive the same licking actions, while selectivity related to sensory stimulus and movement execution remained surprisingly stable. At the same time, the previously acquired motor memories were retained: re-learning to make the same licking actions under the previous context re-activated the previous preparatory activity, even months later. Across learning multiple task contexts, multiple preparatory states were created to encode the same licking action in a context-dependent manner. Our results show that preparatory activity reflects motor memories that stably encode learned actions in combination with their context, which we call a combinatorial code. A feedforward network that stored sensorimotor combinations in high-dimensional hidden layers was able to explain multiple aspects of the results.

Context-specific motor memories may help reduce interference of new learning to previously learned representations ^26,60^, thus protecting existing motor repertoire from erasure in the face of continual learning.

### A paradigm for continual learning that requires motor cortex

Studying how learned actions are stored in motor memory requires a paradigm that can continuously track neural representation of the same movement over extended time. A challenge is that movement kinematics can change over time, obscuring the relationship between the changing neural activity and behavior. Moreover, most existing paradigms test motor learning in single tasks, leaving unclear how motor memories are formed and maintained across continual learning of new motor skills.

To address these challenges, we studied a stereotyped and yet cortex-dependent movement, goal-directed directional licking in mice ^63,64^. We developed a home-cage system in which mice can voluntarily engage in head-fixation and continuously learn multiple licking tasks without human supervision ^65^ (Fig. 1c). In a tactile instructed licking task, mice discriminated the location of a pole (anterior or posterior) presented during a sample epoch and reported their decision using directional licking (“lick left” or “lick right”) after a 1.3 s delay epoch. An auditory ‘go’ cue signaled the motor response (Fig. 1d). Mice initially learned to lick left for anterior pole position and lick right for posterior pole position (task context 1, Fig. 1d). After mice consistently maintained a high level of behavioral performance (>75% correct; Methods), the home-cage system automatically introduced a task rule switch (‘contingency reversal’), altering the association between pole locations and lick directions (task context 2, Fig. 1d). Importantly, the delay epoch separated sensory stimuli from the motor response in time. Thus in the two tasks, mice made identical actions under identical external environment after the delay epoch, but with different stimulus history and task rules. We therefore refer to these conditions as different ‘task context’.

Mice continuously learned many rounds of reversals in the automated home-cage over several months (Fig. 1e). High-speed videography (Methods) showed that the tongue and jaw movements were highly consistent over extended training time and across contingency reversals (Extended Data Fig. 1a-b). Learning as measured by time to reach criterion performance (>75% correct) was faster for subsequent reversals (Fig. 1f; Methods). Faster reversal was observed when mice re-learned the previously learned sensorimotor contingency, but less correlated with overall amount of prior training (a learning to learning effect, Extended Data Fig. 1c), consistent with a saving effect typically associated with motor skill learning ^66^.

The anterior lateral motor cortex (ALM) is critical for planning and execution of directional licking ^19,61,64,67^, but it is unclear whether ALM is continuously required for learned directional licking after extended training. For example, the motor cortex might become disengaged from controlling movement after motor skill acquisition ^68,69^. To test this, we combined the automated home-cage platform with an optogenetic approach to silence ALM activity during task performance in home-cage ^65^ (Methods; Fig. 1g). We virally expressed a red-shifted channelrhodopsin (ChRmine ^70^) in ALM GABAergic neurons and used red light (633 nm) to photostimulate ALM through a clear skull implant during voluntary head-fixation (Fig. 1g). In highly trained mice, photoinhibition of ALM during the delay epoch consistently disrupted behavioral performance, even after multiple rounds of contingency reversal learning (Fig. 1h). Left ALM photoinhibition biased future licking to the ipsilateral direction (lick left) in a light dose dependent manner (Fig. 1i and Extended Data Fig. 1d). These results show that directional licking consistently depends on ALM preparatory activity over time, thus allowing us to chronically track neural activity causally driving the learned licking actions.

### Stable representation of action within the same task context

Do neural representations of learned motor actions remain stable or drift over time (Fig. 2a)? To examine this, we performed longitudinal two-photon calcium imaging of ALM excitatory neurons (GP4.3 mice, Methods; Extended Data Fig. 1e-g; total imaging duration across all experiments, 26-233 days). After mice attained high levels of performance under task context 1 in the home-cage, we transferred the mice to an imaging setup where they performed the same task in daily sessions under a two-photon microscope (Methods). By utilizing an array of over twenty home-cages running in parallel ^65^, this platform enabled us to image trained mice with sufficient throughput while other mice were trained concurrently. After brief acclimation to the imaging setup, mice maintained stable task performance throughout the imaging sessions (Fig. 2b), with little performance change within session (Extended Data Fig. 1j). We imaged the same field of view over ALM across multiple days (Fig. 2c and Extended Data Fig. 2a; referred to as ‘expert-early’ or ‘expert-late’ sessions), covering different fields of view on interleaved days (Extended Data Fig. 2b). We identified neurons that could be matched with high confidence across days based on their centroid locations and the spatial regions contributing to their fluorescence ^71^ (Methods; Extended Data Fig. 2c-d). The imaged fields of view were remarkably stable, yielding 42,739 matched neurons (Extended Data Fig. 2e-i; 50 fields of view, 8 mice).

**Figure 2.**
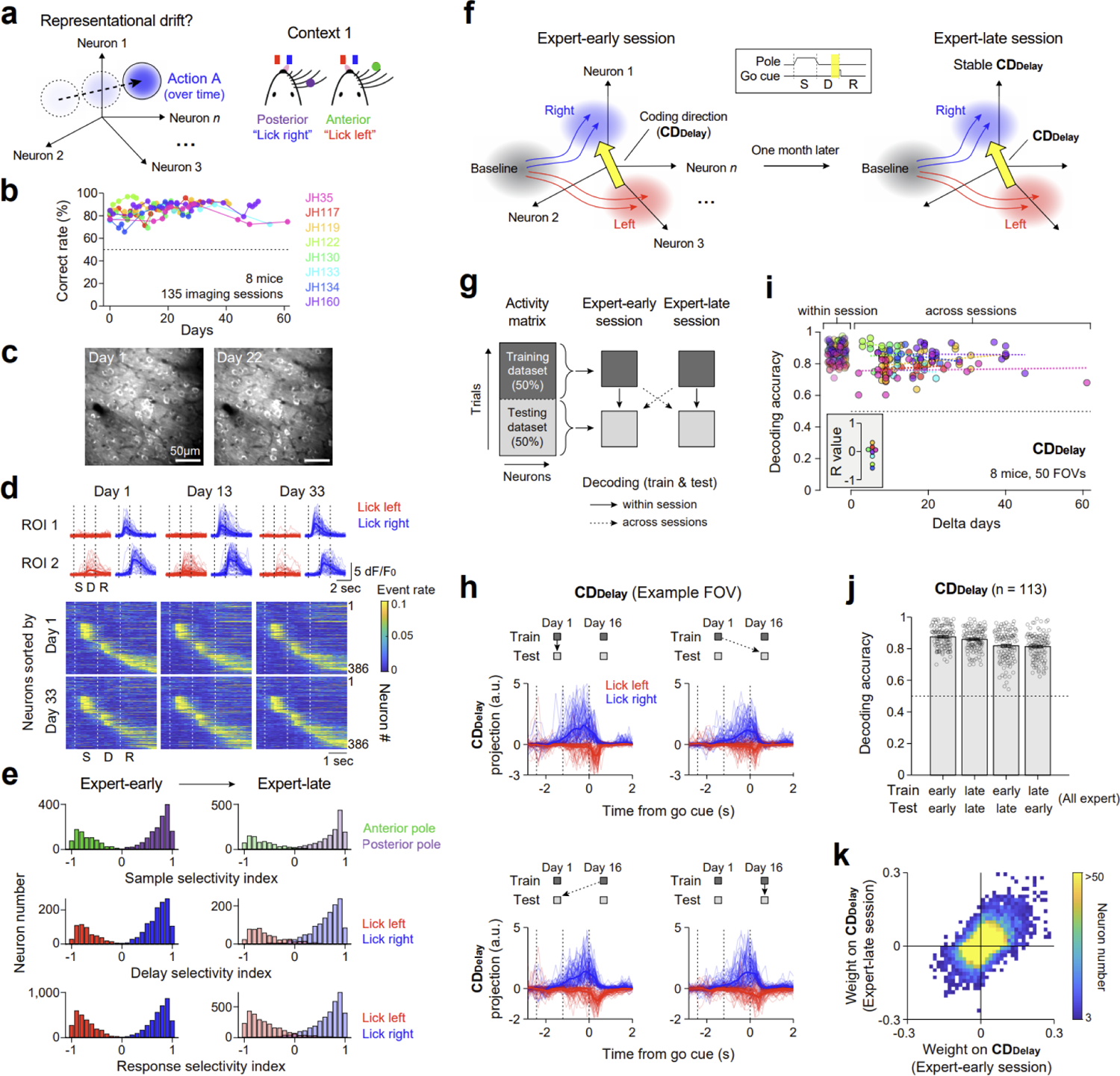
Stable task-related activity over time within the same task context. a. Left, possible outcomes of preparatory states for the same action over time. Right, task context 1. b. Behavior performance during imaging sessions. Each data point shows average performance in one imaging session. Colors indicate individual mice. c. Example images from the same field of view across 2 imaging sessions (Day 1 and 22) showing single neurons expressing GCaMP6s. d. Top, dF/F_0_ activities from two example neurons across days. Thick lines represent the mean; thin lines represent individual trials. Bottom, mean deconvolved activities of all neurons (n=386) registered across days from an example field of view. Neurons are sorted based on their peak activities from different days. e. Histogram of selectivity index in expert-early (left) and expert-late (right) sessions from the same neuronal populations showing statistically significant trial-type selectivity (P<0.001, two-tailed t-test) during the sample (top), delay (middle), and response epoch (bottom) in the expert-early session. Green, neurons preferring anterior pole position trials. Purple, neurons preferring posterior pole position. Red, neurons preferring lick left. Blue, neurons preferring lick right. Trial-type preferences are determined in the expert-early session. f. Schematic of movement-specific activity trajectories in activity space. Coding direction (**CD_Delay_**) is estimated using neural activities during the late delay epoch (inset, yellow shade). Red shade, preparatory state for lick left. Blue shade, preparatory state for lick right. g. Decoding scheme. We used non-overlapping trials for training and testing within (solid arrows) and across imaging sessions (dotted arrows). h. Single-trial ALM activities from an example field of view projected on the **CDDelay** from day 1 (top) or day 16 (bottom). Top left and bottom right plots are projections within session (solid arrows), whereas top right and bottom left plots are projections across sessions (dotted arrows). Thick lines represent the mean; thin lines represent single trials. i. Lick direction decoding using the **CDDelay** as a function of delta days between imaging sessions. Circles with different colors represent different mice. Within session decoding accuracy is the mean of two conditions (train expert-early and test expert-early session, train expert-late and test expert-late session, solid arrows in **g**). Across sessions decoding accuracy is the mean of two conditions (train expert-early and test expert-late session, train expert-late and test expert-early session, dotted arrows in **g**). Dotted lines indicate linear regressions of individual mice across days. Inset, R values of linear regressions of individual mice. j. Decoding accuracy of the **CD_Delay_** within and across imaging sessions under the same task context. n=113 pairs of sessions, 10 mice; mean ± s.e.m. k. Weight contributions of individual neurons to the **CD_Delay_**’s from expert-early and expert-late imaging sessions. 35,420 neurons from 8 mice.

Individual ALM neurons exhibited task-related activity (dF/F_0_, Fig. 2d, top). For further analyses, we deconvolved dF/F_0_ activity to avoid the spillover influence of slow-decaying calcium dynamics across task epochs ^72^ (Methods; Extended Data Fig. 2j). Sorting the same neuronal population based on their peak activities from different days showed similar patterns of task-related activity across days (Fig. 2d, bottom). We computed neuronal selectivity as the difference in activity between trial types divided by their sum (anterior versus posterior pole position for the sample epoch; lick left versus lick right for the delay and response epochs; correct trials only, Methods). On error trials, when mice licked in the opposite direction to the instruction provided by pole location, ALM activity during the delay epoch predicted the licking direction (Extended Data Fig. 2k-l). Neurons showing significant trial-type selectivity (P<0.001, two-tailed t-tests) in expert-early sessions largely maintained their selectivity in expert-late sessions (Fig. 2e; Pearson’s correlation, expert-early vs. expert-late selectivity index, sample epoch: R=0.9404, P=0; delay epoch: R=0.8861, P=0; response epoch: R=0.9001, P=0). Interestingly, a subset of ALM neurons exhibited altered task-related activity across days, but these changes mainly occurred in non-selective neurons (Extended Data Fig. 3a-c). This suggests that lick direction encoding of ALM preparatory activity may be selectively maintained over time.

To investigate the stability of lick direction encoding at the population level, we analyzed ALM activity in an activity space, where each dimension corresponds to the activity of an individual neuron ^19,73^. We estimated a ‘coding direction’ (**CD_Delay_**) in activity space along which ALM activity maximally discriminated future lick direction at the end of the delay epoch (‘preparatory state’, Methods). In the same neuronal population, the **CD_Delay_** allowed us to examine whether ALM activity encoding future lick direction was maintained or reorganized over time (Fig. 2f). We estimated the **CD_Delay_** using ALM activity on 50% of the trials in a session (training dataset) and examined activity projections in non-overlapping trials from the same session or across sessions (testing dataset, Fig. 2g). ALM activity projected on the **CD_Delay_** was maintained over time (Fig. 2h), despite moderate changes in population activity vector (Extended Data Fig. 3d-f). We used a decision boundary on the **CD_Delay_** to predict lick left versus lick right from ALM activity (Methods). A **CD_Delay_** decoder defined in one session could be used to accurately predict lick direction in other sessions regardless of the timespan between sessions, even up to 2 months apart (Fig. 2i), with no detectable decrease in decoding performance over time (linear regression: −0.08 ± 0.11, mean ± s.e.m. across mice; P=0.4870, t-test against 0). The **CD_Delay_** decoders defined in expert-early or late sessions could similarly predict lick direction when applied to expert-late or early sessions, respectively (Fig. 2j). Moreover, individual neurons contributing to the **CD_Delay_** were highly correlated across sessions (Fig. 2k; Pearson’s correlation, R=0.6053, P=0).

We applied the same analyses to ALM activity during the sample and response epochs and found similarly stable selectivity along the coding directions (Extended Data Fig. 4). These results show that, in the same task context, ALM activity is selectively maintained along coding directions that encode learned directional licking for at least 2 months.

### New representation emerges with learning of new task context

How do motor memories form when we acquire new motor skills? Are existing activity states reused ^9,46^ or do entirely new activity states form under new context (Fig. 3a)? To address this question, we monitored activity in the same ALM population across two different task contexts. After imaging ALM activity in task context 1, we returned mice to the home-cage system where they learned the reversed sensorimotor contingency (Extended Data Fig. 1f). We then transferred them to the two-photon setup again and imaged the same fields of view in task context 2 (Fig. 3b; task context 1→2).

**Figure 3.**
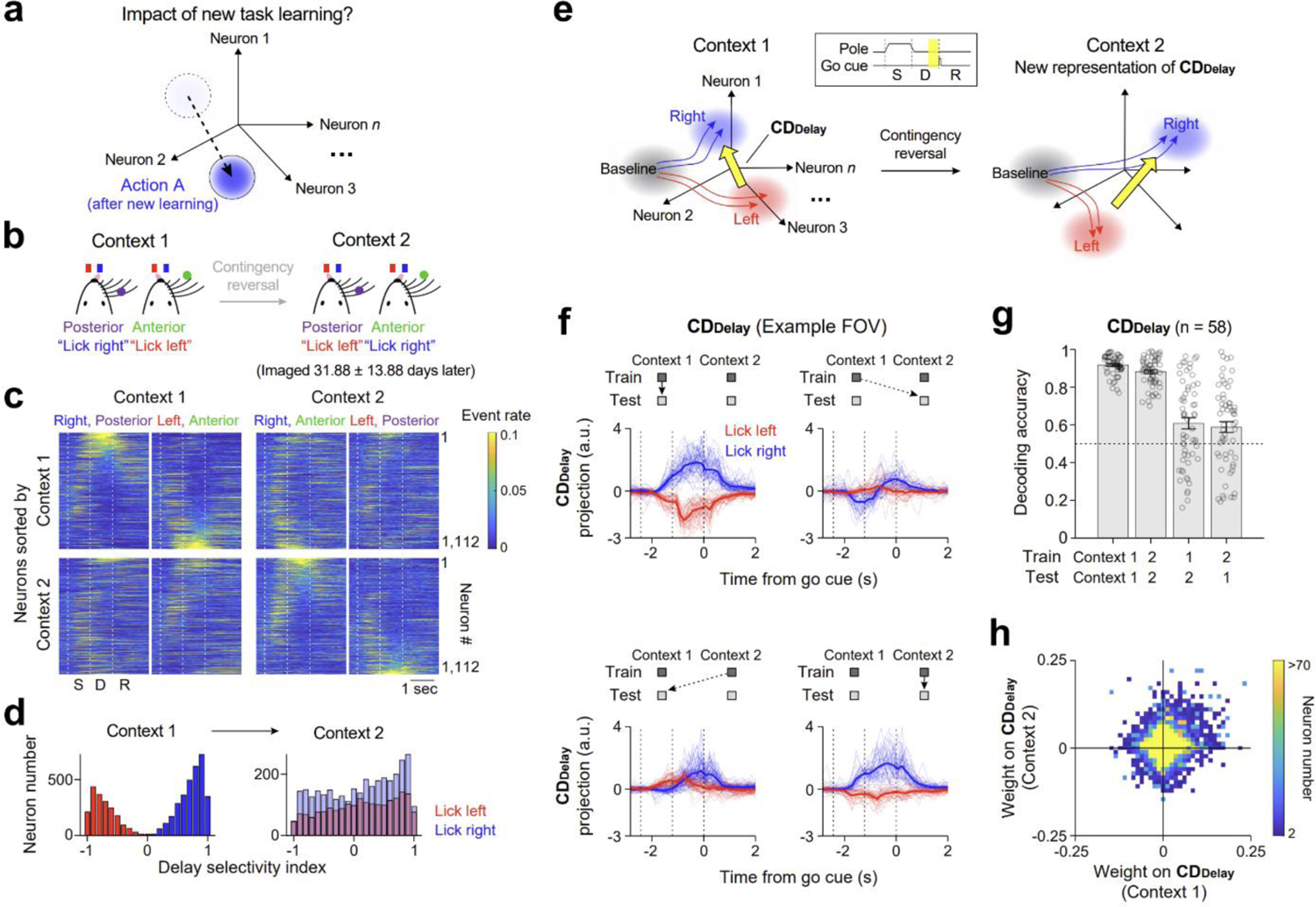
New preparatory activity emerges by learning new task context. a. Possible outcomes of preparatory states across different task contexts. b. Task structures of task context 1 (left) and 2 (right). Time interval between imaging sessions is 31.88 ± 13.88 days, mean ± SD across fields of view. c. Mean deconvolved activities from an example field of view across task contexts (n=1,112 neurons). Neurons are sorted based on their selectivity amplitude during the delay epoch in either task context 1 (top row) or task context 2 (bottom row). d. Histogram of selectivity index in task context 1 (left) and 2 (right) from the same neuronal populations showing statistically significant trial-type selectivity (P<0.001, two-tailed t-test) during the delay epoch in task context 1. Red, neurons preferring lick left. Blue, neurons preferring lick right. Trial-type preferences are determined in task context 1. e. Schematic of movement-specific activity trajectories in activity space and **CD_Delay_**’s across task contexts. Red shade, preparatory state for lick left. Blue shade, preparatory state for lick right. f. Single-trial ALM activities from an example field of view projected on the **CDDelay** from task context 1 (top) or task context 2 (bottom). Top left and bottom right plots are projections within the same task context (solid arrows), whereas top right and bottom left plots are projections across different task contexts (dotted arrows). Thick lines represent the mean; thin lines represent single trials. g. Decoding accuracy of the **CD_Delay_** within and across task contexts. n=58 pairs of sessions, 10 mice; mean ± s.e.m. Individual circles show individual fields of view. h. Weight contribution of individual neurons to the **CD_Delay_**’s from task contexts 1 and 2. 44,409 neurons from 10 mice.

Contingency reversal allowed us to specifically alter the task context while keeping the sensory instruction and motor response identical, thereby avoiding neural activity changes caused by changes in movement parameters. Mice showed consistent task performance in the imaging sessions across the two task contexts (85.59 ± 1.00% vs. 84.06 ± 0.99% correct rate, mean ± s.e.m; P=0.1862, paired t-test). Video analysis showed that the tongue and jaw movements were indistinguishable across the two task contexts, confirming that mice made the same actions under the two conditions (Extended Data Figs. 1a-b, bottom). We identified matched neurons across imaging sessions, yielding an average of 1,118 ± 500 neurons in each field of view (58 fields of view, 10 mice; time gap between task contexts 1 and 2 imaging sessions, 31.88 ± 13.88 days, mean ± SD across sessions).

We observed a profound reorganization of ALM preparatory activity after mice learned new task context. Many ALM neurons showing prominent lick left or lick right preferring activities during the delay epoch in task context 1 lost or even reversed their selectivity in task context 2 (Fig. 3c, top), while other neurons retained their selectivity. At the same time, new selective neurons emerged in task context 2 (Fig. 3c, bottom). Across the population, neurons with significant trial-type selectivity during the delay epoch in task context 1 showed nearly random selectivity distribution in task context 2 (Fig. 3d and Extended Data Fig. 5e; Pearson’s correlation, task context 1 vs. 2 selectivity index, R=-0.0057, P=0.6774).

We examined the encoding of future lick direction in the same ALM population by calculating the **CD_Delay_** in each task context (Fig. 3e). Single-trial activity projected on the **CD_Delay_** reliably differentiated lick left and lick right within task context, but this separation largely collapsed when projected on the **CD_Delay_** across task contexts (Fig. 3f). Across all fields of view, a **CD_Delay_** decoder trained to predict lick direction in one task context performed at near chance level on average in the other task context (Fig. 3g). Additionally, individual neurons supporting the **CD_Delay_**’s in the two task contexts were weakly correlated (Fig. 3h; Pearson’s correlation, R=0.3; significantly less than the Pearson’s correlation within task context over time in Fig. 2k, P=0, bootstrap), consistent with the reshuffling of single neuron selectivity (Fig. 3d and Extended Data Fig. 5e). Thus, task contexts 1 and 2 yielded distinct **CD_Delay_**’s. In contrast to the profound reorganization of ALM preparatory activity, selectivity during the sample and response epochs remained remarkably stable across task contexts (Extended Data Fig. 5). This ruled out the possibility that the observed reorganization of preparatory activity was due to unstable imaging over time, or changes in the mice’s motor behavior.

Although ALM preparatory activity was reorganized across task contexts on average, we found substantial individual variability across mice (Fig. 3g and Extended Data Fig. 6a-c). In some mice, the **CD_Delay_** in the new task context was near orthogonal to the **CD_Delay_** from the previous context in activity space (Fig. 3f). But in other mice, preparatory activity maintained along the previous **CD_Delay_** (Extended Data Fig. 6d), or even reversed direction along the previous **CD_Delay_** (Extended Data Fig. 6e). Within each mouse, the similar pattern of reorganization was consistently observed across different imaging fields of view (Extended Data Fig. 6a-c), indicating that the variability was not due to heterogeneous sampling of neurons, variable estimates from the finite number of trials, or spatial location of imaging (Extended Data Fig. 6g). This individual variability was also not explained by task performance, ongoing movements during the delay epoch, the speed of task learning, or the time intervals between the task contexts (Extended Data Fig. 6f-g). Individual variability may result from differences in the underlying circuits (see later modeling results).

These results show that new preparatory states form when mice learn to make the same licking actions under new task context. These results also show that there is no unique preparatory state for specific movements. Distinct preparatory states in the motor cortex can drive the same subsequent movement execution. This suggests that the preparatory states could encode a learned action in multiple representations that index distinct contexts.

### Stable retention and re-activation of learned representations

If learned actions are encoded in combination with their contexts, this could enable stable retention of motor memories over continual learning, because new learning in different contexts forms parallel new representations without altering the previously learned representations. To test this notion, we re-examined ALM activity in the previous context after the intervening learning. If preparatory states reflect a consistent motor memory of learned actions and their contexts, generating the same actions in the previous context should re-activate the previously learned preparatory states (Fig. 4a).

**Figure 4.**
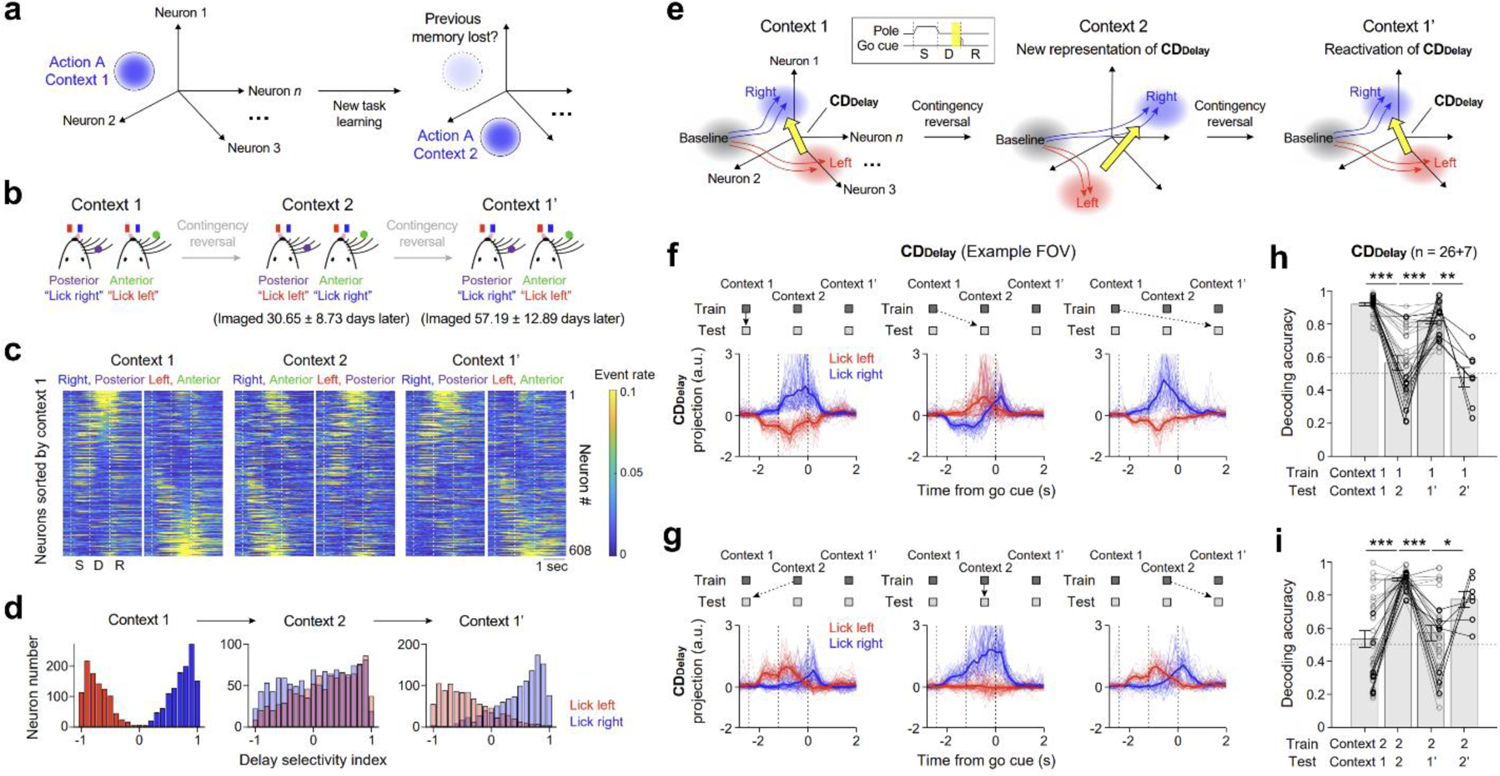
Re-learning previous task context re-activates the learned preparatory activity. a. Possible outcomes of preparatory states over the course of re-learning (task 121’). b. Task structure of context 1 (left), context 2 (middle), and context 1’ (right). Task contexts 1 and 1’ are identical but mice experienced intervening learning in task context 2. Time interval between imaging in task contexts 1 and 2 is 30.65 ± 8.73 days and 57.19 ± 12.89 days between task contexts 1 and 1’, mean ± SD across fields of view. c. Mean deconvolved activities from an example field of view across three task contexts (n=608 neurons). Neurons are sorted based on their selectivity amplitude during the delay epoch from task context 1. d. Histogram of selectivity index in task context 1 (left), 2 (middle), and 1’ (right) from the same neuronal populations showing statistically significant trial-type selectivity (P<0.001, two-tailed t-test) during the delay epoch in task context 1. Red, neurons preferring lick left. Blue, neurons preferring lick right. Trial-type preferences are determined in task context 1. e. Schematic of movement-specific activity trajectories in activity space and **CD_Delay_**’s across task contexts. Red shade, preparatory state for lick left. Blue shade, preparatory state for lick right. f. Single-trial ALM activities from an example field of view projected on the **CDDelay** from task context 1. Projections in task context 1 (left), task context 2 (middle), or task context 1’ (right). Thick lines represent the mean; thin lines represent single trials. g. Same as **f**, but for the **CD_Delay_** from task context 2. h. Decoding accuracy of the **CD_Delay_**’s from task context 1 tested on task contexts 1, 2, 1’, and 2’. Gray circles and lines indicate fields of views imaged across task contexts 1, 2, and 1’ (n=26 fields of view, 5 mice). Black circles and lines indicate a subset of fields of views imaged across task contexts 1, 2, 1’, and 2’ (n=7 fields of view, 3 mice). Task context 1 vs. 2, ***P=1.12×10^-10^; task context 2 vs. 1’, ***P=3.46×10^-7^; task context 1’ vs. 2’, **P=0.0091. Paired t-test. i. Same as **h**, but for the **CD_Delay_**’s from task context 2. Task context 1 vs. 2, ***P=4.87×10^-10^; task context 2 vs. 1’, ***P=7.40×10^-9^; task context 1’ vs. 2’, *P=0.0496. Paired t-test.

After imaging ALM activity in task contexts 1 and 2, mice were re-trained in the previous task context 1 (notated as 1’ for re-learning) in the automated home-cage (Extended Data Fig. 1f). We then imaged neuronal activity from the same ALM population across task contexts (Fig. 4b; task context 1→2→1’). We observed a re-activation of the previous preparatory activity pattern after mice re-learned the previous task, even though task contexts 1 and 1’ were tested 2 months apart on average (range: 32-78 days across mice; Fig. 4b and Extended Data Fig. 1f-g). Individual ALM neurons showing lick left or lick right preferring activity in task context 1 were reconfigured in task context 2 but reappeared in task context 1’ (Fig. 4c). Across the population, neurons with significant trial-type selectivity during the delay epoch in task context 1 showed nearly random selectivity distribution in task context 2, but upon re-testing in task context 1’, a similar distribution of selectivity re-emerged (Fig. 4d and Extended Data Fig. 7h; Pearson’s correlation, task context 1 vs. 1’ selectivity index, R=0.7675, P=0).

We examined whether ALM population encoding of future lick direction was similarly re-activated in task context 1’ (Fig. 4e). Activity trajectories in lick left and lick right trials were well separated in task context 1’ when projected on the **CD_Delay_** from task context 1 (Fig. 4f), indicating a re-activation of preparatory activity along similar **CD_Delay_**’s in activity space. In contrast, the activity trajectories were poorly separated when projected on the **CD_Delay_** from task context 2 (Fig. 4g), consistent with a reorganization of preparatory activity across task contexts. Across all fields of view, a **CD_Delay_** decoder trained on task context 1 predicted lick direction at near chance level on average in task context 2, but its performance recovered in task context 1’ (Fig. 4h). Together, these data indicate a re-activation of the previously learned preparatory states under task context 1’.

We also observed a similar re-activation of ALM preparatory states associated with task context 2 when mice were re-tested in this task. In a subset of mice (n=3), we further imaged the same ALM populations following one more contingency reversal (task context 1→2→1’→2’, spanning up to 3 months, range: 59-97 days across mice; Extended Data Fig. 1f-g). We found consistent reorganization and re-activation of preparatory activity across the reversals (Fig. 4i). Thus, stable retention of preparatory states was not limited to any one specific task context. Unlike preparatory activity, selectivity during the sample and response epochs were stably maintained across the entire time regardless of task contexts (Extended Data Fig. 7).

In addition to the reorganization and re-activation of ALM preparatory activity encoding lick direction (**CD_Delay_**), we also observed activity changes along other dimensions of activity space across task contexts (Extended Data Fig. 8). Activity along these dimensions did not discriminate lick direction (Extended Data Fig. 8e; ‘movement-irrelevant subspace’). Activity was shifted along the movement-irrelevant subspace after learning new task contexts but did not recover in task context 1’ or 2’ (Extended Data Fig. 8c-d). Therefore, preparatory activity is selectively maintained along coding directions encoding behavior-related information, but activity drifts over time along other non-informative directions ^11,19,74^.

Taken together, these results show that motor learning produces context-specific preparatory states. Once learned, these activity states are stably stored and can be recalled after several months, even despite intervening motor learning involving the same actions in other contexts. These preparatory states thus reflect context-specific motor memories that are stably retained over continual learning.

### Learning creates parallel representations of the same action across contexts

We next asked whether continual learning in new task contexts will continue to create new preparatory states for learned actions. Our experiments thus far examined only two task contexts. Furthermore, contingency reversal may constitute a special case of task context change, which requires a re-association of previously learned sensory stimuli and motor actions. If mice learned to perform directional licking instructed by a novel stimulus, would yet new preparatory states emerge under new task context (Fig. 5a)?

**Figure 5.**
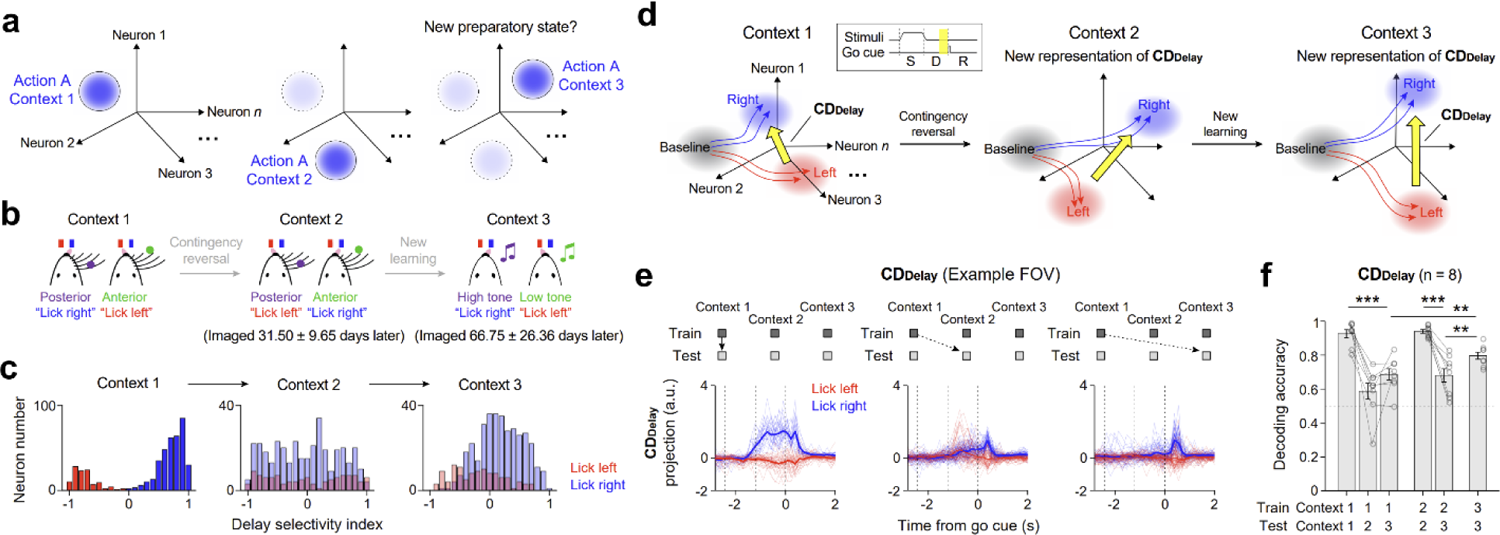
Learning new tasks with novel stimulus creates new preparatory activity. a. Possible outcomes of preparatory states over the course of continual learning (task 123). b. Task structure of task context 1 (left), 2 (middle), and 3 (right). In task context 3, mice discriminate the frequency of a pure tone during the sample epoch and report frequency using directional licking after the delay epoch. Time interval between imaging in task contexts 1 and 2 is 31.50 ± 9.65 days and 66.75 ± 26.36 days between task contexts 1 and 3, mean ± SD across fields of view. c. Histogram of selectivity index in task context 1 (left), 2 (middle), and 3 (right) from the same neuronal populations showing statistically significant trial-type selectivity (P<0.001, two-tailed t-test) during the delay epoch in task context 1. Red, neurons preferring lick left. Blue, neurons preferring lick right. Trial-type preferences are determined in task context 1. d. Schematic of movement-specific activity trajectories in activity space and **CD_Delay_**’s across task contexts. Red shade, preparatory state for lick left. Blue shade, preparatory state for lick right. e. Single-trial ALM activities from an example field of view projected on the **CDDelay** from task context 1. Projections in task context 1 (left), task context 2 (middle), or task context 3 (right). Thick lines represent the mean; thin lines represent single trials. f. Decoding accuracy of the **CD_Delay_**’s from task context 1 tested on task contexts 1, 2, and 3 (n=8 fields of view, 3 mice). Task context 1 vs. 3, ***P=5.56×10^-5^. Decoding accuracy of the **CD_Delay_**’s from task context 2 (n=8 fields of view, 3 mice). Task context 2 vs. 3, ***P=6.08×10^-4^. Decoding accuracy of the **CD_Delay_**’s from task context 3 (n=8 fields of view, 3 mice). Comparing to task context 1 decoder, **P=0.0022; comparing to task context 2 decoder, **P=0.002. Paired t-test.

To address this question, we trained mice to perform an auditory instructed directional licking task (task context 3) in the automated home-cage after imaging ALM activity in the tactile tasks (Fig. 5b; task context 1→2→3; range: 40-118 days across mice). In this auditory task, mice learned to perform directional licking based on the frequency of a pure tone presented during the sample epoch (lick left for 2 kHz, lick right for 10 kHz).

After reaching criterion performance, we imaged the same ALM populations and compared neuronal activity across the three task contexts. Individual ALM neurons with significant selectivity for lick left or lick right during the delay epoch in tactile task context 1 showed distinct pattern of selectivity in the auditory task (task context 3) as well as in tactile task context 2 (Fig. 5c; Pearson’s correlation, task context 1 vs. 3 selectivity index, R=0.3435; significantly less than the Pearson’s correlation within task context over time in Fig. 2e, P=0, bootstrap).

We further examined if ALM preparatory activity in the tactile and auditory tasks encoded future lick direction along different dimensions in activity space (Fig. 5d, **CD_Delay_**). Indeed, we found poor separation between single-trial activity trajectories in lick left and lick right trials when ALM activities in the auditory task were projected on the **CD_Delay_** from the tactile task (Fig. 5e). Across all fields of views, the **CD_Delay_** decoders trained on the tactile tasks (either task context 1 or 2) exhibited poor performance in predicting lick direction when tested on the auditory task (Fig. 5f). In contrast, a decoder trained within the auditory task could decode lick direction significantly better than the decoders trained on the tactile tasks (auditory task decoder vs. tactile task 1 decoder and tactile task 2 decoder, P=0.0022 and P=0.002, two-tailed paired t-test), indicating that their poor decoding performances in the auditory task were not due to a lack of neuronal selectivity.

Finally, ALM activity during the sample epoch was distinct across tactile and auditory tasks, which was expected given the distinct sensory inputs (Extended Data Fig. 9a-b). Notably, lick direction selectivity during the response epoch remained stable across all three task contexts, which likely reflected conserved licking movement execution across tasks (Extended Data Fig. 9e-f). Stable selectivity during the response epoch also ruled out the possibility that the observed reorganization of preparatory activity was due to unstable imaging over time.

These results show that motor learning in new task contexts keeps creating new preparatory states for the same action in a context-dependent manner. At the same time, activity related to movement execution remains the same across contexts. Thus, learned actions are encoded by distinct preparatory states in combination with their contexts.

### Preparatory activity reflects motor memory

How does a context-specific neural code support motor memory behavior? Mice exhibit faster re-learning in the previously learned sensorimotor contingency (Extended Data Fig. 1c). Do ALM preparatory states retain a memory trace of the previous learning ^26^? Preparatory states in different task contexts are arranged along distinct coding directions in activity space. We examined if each task-specific coding direction retained a memory of the previous learning in the same task.

We re-analyzed the chronological two-photon imaging dataset from the tactile tasks (task context 1→2→1’) in which we imaged ALM activity in the same task context before and after an intervening learning. If learning of the task context 2 left a memory trace in ALM activity, we should observe an activity change in task context 1’ compared to task context 1, and this change should support the performance of task 2. To look for such change, we calculated the **CD_Delay_** for task context 2 and projected ALM activity at the end of the delay epoch on the **CD_Delay_** (Extended Data Fig. 10a). After learning in task context 2, ALM activity in task context 1’ exhibited increased lick direction selectivity along the **CD_Delay_** compared to task context 1 (Extended Data Fig. 10b; P=0.005, paired t-test). To examine if this activity change could support the performance of task 2, we performed decoding of lick direction using activity projected on the **CD_Delay_** from task context 2. Decoding was near chance level in task context 1 (52.75 ± 5.24%, mean ± s.e.m. across sessions) but increased to 58.66 ± 4.63% in task context 1’ (Extended Data Fig. 10c; P=0.0199, paired t-test). This shows that learning of task context 2 left a subtle but persistent alteration of ALM preparatory activity along the **CD_Delay_** that retained a memory trace of the previous learning ^26^.

If each task-specific **CD_Delay_** retains a memory trace of the previous learning, the **CD_Delay_**’s could provide a place to store task-specific motor memories without interference. We examined whether the ability to create context-specific coding directions could protect existing memories. We took advantage of the individual variability across mice and tested whether mice with reorganized **CD_Delay_**’s across task contexts exhibited faster re-learning (i.e. greater saving effect) than mice with fixed **CD_Delay_**’s (Extended Data Fig. 6a-c). Remarkably, the degree of **CD_Delay_** reorganization across task contexts 1 and 2 predicted the mice’s re-learning speed in task context 1’ (Extended Data Fig. 10d, P=0.0002, Pearson’s correlation). If a mouse exhibited more distinct **CD_Delay_**’s across different task contexts (lower dot product), the mouse re-learned the previously learned task faster.

These results suggest that task-specific motor memories are stored along distinct coding directions in activity space, which could help protect the memories from interference by new learning and support faster re-learning of the previously learned tasks.

### Stable memory storage is consistent with a feedforward network architecture

We next used network modeling to explore network architectures that might support the observed memory storage. Preparatory activity is mediated by interactions between ALM and multiple brain regions ^13^. Our goal was to explore what networks could explain how preparatory activity becomes reorganized by learning while being agnostic to how the models map onto brain regions. Specifically, we focused two key features of the neural results: 1) formation of new preparatory activity patterns during contingency reversal learning; 2) stable retention of learned preparatory activity patterns that can be re-activated after intervening task learning.

We started with recurrent neural networks (RNNs) due to their ability to discover models that are constrained to solve a task without enforcing their solutions ^75^ (Fig. 6a). RNNs learned to generate choice-specific persistent activity after a transient input (Fig. 6b, task context 1). To examine contingency reversal learning, we trained the internal connections of the learned RNNs to generate the opposite responses (lick direction) to the previous inputs (task context 2) while keeping the input and output connections fixed (Methods). Notably, this standard RNN architecture was unable to capture all features of the neural data. After reversal learning, the selectivity of the RNN units mostly followed the network output (i.e. lick direction, Fig. 6b). Network units contributing to the **CD_Delay_** (defined by lick direction) in task context 1 similarly contributed to the **CD_Delay_** in task context 2 (Fig. 6c, task context 1 vs. 2). To rule out the possibility that the network output and learning rule dictated the dynamics of the internal units, we also tested RNNs in which only two internal units contributed to the output, yielding similar results (Extended Data Fig. 11a-c). The RNN dynamics were therefore constrained to one dimension along the previously learned **CD_Delay_** and the networks solved the contingency reversal by re-association (Fig. 6d). This is contrary to the neural data in which a new pattern of selectivity emerged during contingency reversal learning (Fig. 3).

**Figure 6.**
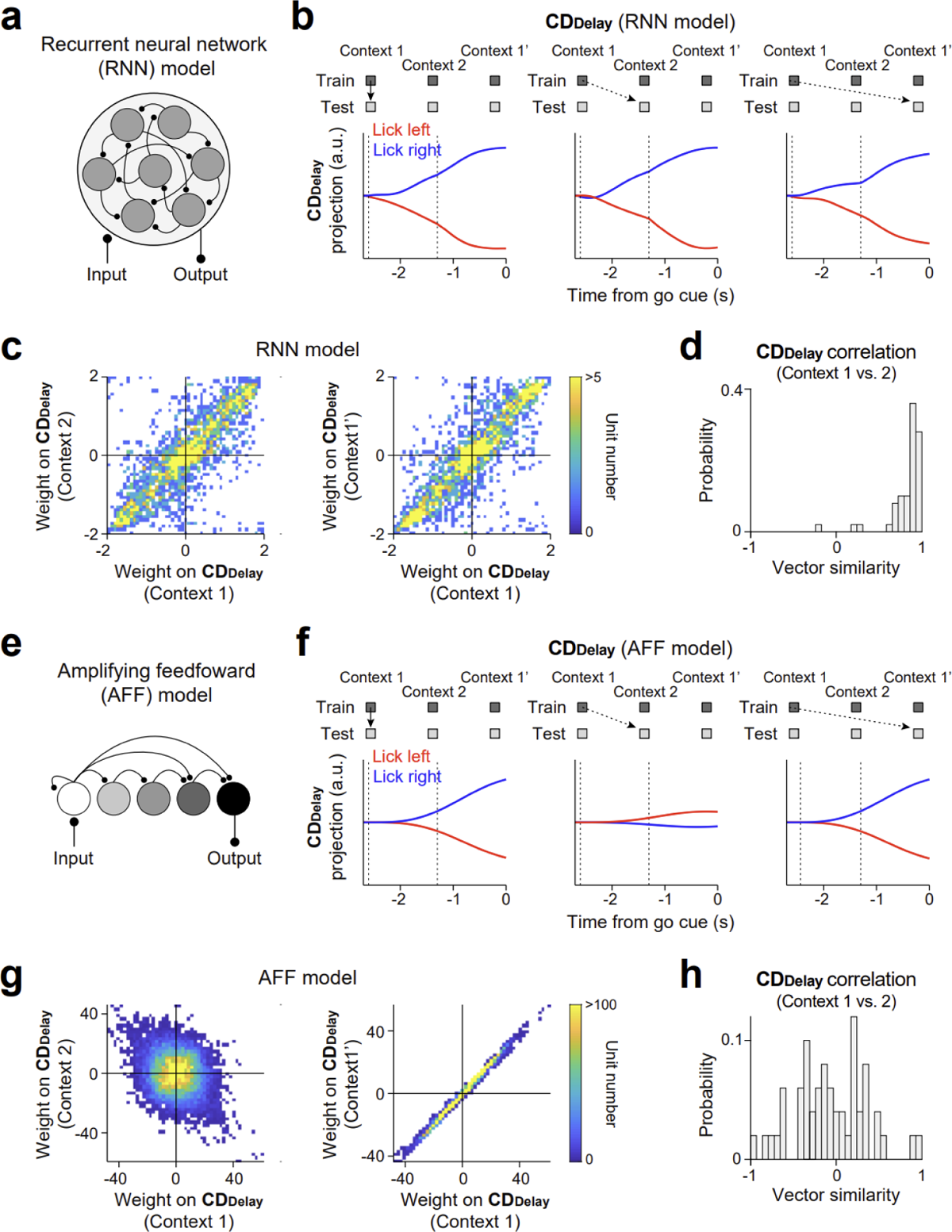
Neural network modeling. a. Schematic of the recurrent neural network (RNN) model. b. Activity of a RNN model projected on the **CDDelay** from task context 1. Projections in task context 1 (left), task context 2 (middle), or task context 1’ (right). The **CDDelay** is defined by lick direction. Blue, lick right. Red, lick left. c. Weight contribution of the RNN units to the **CD_Delay_**’s from task contexts 1 and 2 (left), or the **CD_Delay_**’s from task contexts 1 and 1’ (right). d. Dot product between the **CD_Delay_**’s from the RNNs across task contexts 1 and 2. Data from 50 randomly initialized RNNs. e. Schematic of the amplifying feedforward (AFF) model. **f-h**. Same as **b-d**, but for the AFF models. Data from 50 randomly initialized AFFs.

We next explored a class of amplifying feedforward (AFF) networks that generate persistent activity by passing activity through a chain of network states ^76,77^ (Fig. 6e and Extended Data Fig. 11d), which can be modeled as a series of layers with feedforward connections. The AFF networks learned feedforward amplifications to generate choice-specific persistent activity in response to transient inputs to the early layer (Fig. 6f) and can explain a variety of previous findings from ALM perturbation experiments ^77^.

Feedback connections from late layers to early layers conveyed output signals and allowed the network to learn via error backpropagation (Methods). In the hidden layers, the AFF networks maintained persistent activity along multiple dimensions (Extended Data Fig. 11e-f). The AFF networks readily captured both features of the neural data: 1) upon contingency reversal, the network learned a new pattern of selectivity that was distinct from the previous selectivity pattern, i.e., network activity in task context 2 showed collapsed activity on the **CD_Delay_** from task context 1 (Fig. 6f); 2) upon re-training in the previous sensorimotor contingency, the network re-activated the previous pattern of selectivity (Fig. 6f, task context 1’). Resetting the weights of the hidden layers before re-training in task context 1’ prevented the re-activation of the previous **CD_Delay_** (Extended Data Fig. 11g-h) Thus, the AFF networks stored sensorimotor mappings in high-dimensional hidden layers.

We examined the features of the AFF networks that allowed them to create new **CD_Delay_**’s upon contingency reversal learning while stably retaining previously learned **CD_Delay_**’s. Sensory inputs to the early layer of the AFF networks are gradually transformed into appropriate motor outputs in the late layers. Due to feedforward connections (containing stimulus input activity) and feedback connections (containing motor output activity), intermediate layers contained mixtures of the input and output representations. We decompose AFF network activity into distinct modes. AFF networks learned a persistent stimulus mode and an output mode along orthogonal dimensions (Extended Data Fig. 12a). In the activity space, the persistent stimulus mode combined with the output mode to establish the **CD_Delay_** (Extended Data Fig. 12a, bottom row).

Upon contingency reversal, the output mode combined with the new stimulus mode to form a new **CD_Delay_** (Fig. 6g, task context 1 vs. 2; Extended Data Fig. 12a). Reversion to the previous contingency re-activated the original stimulus and output modes, which re-activated the previously **CD_Delay_** (Fig. 6g, task context 1 vs. 1’). In contrast, we found that the persistent stimulus mode was absent in the RNNs, which resulted in **CD_Delay_**’s that were aligned to only the output mode (Extended Data Fig. 12b). This suggests that a high-dimensional circuit that can maintain multiple persistent activity modes is critical to support context-dependent **CD_Delay_** reorganization.

This feature of the AFF networks could also explain variable degree of activity reorganization observed across individual mice during contingency reversal learning (Extended Data Fig. 6a-c). Because intermediate layers contained mixtures of the input and output representations (Extended Data Fig. 12a), individual networks could exhibit a range of reorganization behavior depending on the relative strength of the input and output representations (Fig. 6h). Networks with strong stimulus modes (due to weak feedback connections) exhibited reorganized **CD_Delay_**’s; networks with strong output modes (due to strong feedback connections) exhibited stable **CD_Delay_**’s aligned to the network output (Fig. 6h). Across all models, the strength of the stimulus mode predicted the degree of **CD_Delay_** reorganization across task contexts (Extended Data Fig. 12c).

This suggests an unexpected role of stimulus activity in the formation of motor memory. We tested whether the strength of stimulus activity in ALM could explain the individual variability across mice in our neural data. Remarkably, stimulus activity strength measured in task context 1 predicted whether a mouse would exhibit context-dependent reorganization of **CD_Delay_** across task contexts (Extended Data Fig. 12d). This suggests that the observed variability across mice may arise from individual differences in their underlying neural circuits.

In summary, an AFF network architecture that maintained multiple persistent activity modes to encode sensorimotor combinations in high-dimensional hidden layers could explain multiple aspects of the neural data. These results suggest that stable motor memory is rooted in high-dimensional representations. We note that the AFF network contains feedback connections and therefore is a subclass of RNNs. There may be other network architectures that could also produce these neural dynamics.

## Discussion

Our study reveals a combinatorial neural code that stores learned actions in combination with their contexts. Within a task context, preparatory activity encoding lick direction is stably maintained over multiple months (Fig. 2), and even across intervening motor learning (Fig. 4). Across task contexts, the same action is preceded by distinct patterns of preparatory activity (Fig. 3), while selectivity related to sensory stimulus and movement execution remains remarkably stable over time and across task contexts (Extended Data Figs. 4, 5, 7, and 9). These results suggest that the same action can be encoded by multiple preparatory activity states. This afforded degree of freedom may allow the motor circuits to create parallel representations for the same actions while indexing their contexts. Indeed, we find that new task learning continually creates new preparatory states for learned actions in a context-dependent manner (Fig. 5).

Preparatory states in different task contexts are arranged along distinct coding directions in activity space. Each coding direction retains a memory trace of the previous learning in specific tasks (Extended Data Fig. 10a-c). Context-specific coding directions could help protect existing memories from interference by new learning: mice that exhibit more distinct coding directions across task contexts were faster to re-learn the previously learned task after intervening learning, i.e. greater saving (Extended Data Fig. 10d). These properties of ALM preparatory activity indicate that it reflects the neural substrate of motor memory. Context-specific memory, as we observed in the motor system, may provide a solution for stable memory storage throughout continual learning. Learning in new contexts produces parallel new representations instead of modifying existing representations, thus protecting existing motor memories from erasure by new motor learning ^26,60^.

Motor cortical preparatory activity is thought to provide the initial conditions for subsequent movement execution ^20^. Our results show that preparatory activity is not directly linked to the movement itself but reflects motor memories of learned actions and context ^25^. Reorganization of preparatory activity across task contexts shares similarities with place cells of hippocampus, which encode space and experience within specific contexts and undergo global remapping across distinct contexts ^78^. Preparatory activity encodes learned actions in specific task context, and preparatory activity for the same action undergoes remapping in new task context; yet the previous preparatory activity pattern is retained and can be re-activated in the previous task context, even after several months. Motor learning thus forms modular motor memories for each context.

Once formed, motor memories are remarkably stable, and even after intervening motor learning involving the same actions. Context-specific memory may be a general feature for learning cognitive representations.

Our findings suggest that when movement parameters and task context are controlled, neural representation of actions in motor cortex shows surprisingly little representational drift. Within the same task context, lick direction selectivity is remarkably stable and can be re-activated months later, even across intervening motor learning in new tasks involving the same actions. Interestingly, preparatory activity is selectively maintained along coding directions, but activity drifts over time along other non-informative directions (movement-irrelevant subspace, Extended Data Figs. 3 and 8). Persistent preparatory activity is maintained by recurrent networks in motor cortex and connected brain areas ^13,15,20^. Our findings suggest that motor memories are stored in stable network configurations. Previous studies have reported representational drift in sensory, association, and memory-related brain regions ^51,79–82^. However, little representational drift has been reported in motor areas ^56–58^. Differences in brain areas and behavioral paradigms may explain some differences in these findings.

Previous studies of motor skill learning have focused on training animals to adapt learned actions to novel mechanical environments ^11,36,37,83^ or to perform instructed movements in novel task context ^31,38–42^. Motor cortex activity undergoes systematic reconfiguration during these motor skill learnings. However, movement kinematics and associated forces often change with learning, which accompany changes in neural activity, thus obscuring which behavioral features are retained in motor memory. Recent brain-computer interface (BCI) experiments circumvent this challenge by experimentally controlling the mapping between neural activity and motor output ^9,43–49^. These experiments reveal remarkable malleability in motor cortex activity: over timescale of hours, motor cortex can rapidly reassociate existing activity patterns with different motor outputs ^45,46^; across days, learning can create new patterns of activity ^84^. However, BCI experiments typically do not delineate preparatory activity and activity driving movement execution. Moreover, these experiments typically examine activity across a short time span. Stable long-term recording from the same neuronal population remains a challenge. In rare cases where motor cortex activity is tracked up to one month or beyond ^56–58^, studies have reported stable patterns of activity around movement execution, mirroring our findings of stable selectivity during movement execution over long-term (Extended Data Figs. 4, 5, 7, and 9).

Sun et al and Losey et al recently reported that motor learning induces a persistent change in preparatory activity, which may serve as an index for motor memories ^11,26^. Notably, this persistent change occurs outside of the neural activity subspace encoding specific movements (coding directions), manifesting in a uniform shift of preparatory states in activity space, while the geometry of the activity states encoding specific movements is mostly preserved. Moreover, these studies mostly examine activity change within a session or across a few days, thus the stability of the reorganized activity remains to be determined. Distinctively, here we find that learning new task context induces a dramatic reorganization of the coding directions (Fig. 3), along with changes in movement-irrelevant subspace (Extended Data Fig. 8). By tracking activity over long-term, we find that for the same movement, each task context induces an orthogonal pattern of preparatory activity and continual task learning keeps creating new preparatory activity patterns (Fig. 5). Notably, we also find that, once learned, the preparatory states are stably retained in memory and can be recalled even after multiple months (Fig. 4). Thus multiple concerted changes, along both coding directions and movement-irrelevant subspaces, may accompany learning of new motor skills. These distinct mechanisms may work collectively to increase storage capacity and help differentiate motor memories.

Previous studies of motor learning mostly examine activity within a single task. Here we take advantage of a directional licking behavior and automated home-cage training in mice to track neural representation of learned actions across multiple tasks while conserving movement parameters. Importantly, learned directional licking requires ALM (Fig. 1), which allows us to examine neural activity causally driving the learned actions. Unsupervised home-cage training in parallel (>20 cages) is critical to ensure we can image trained mice with sufficient throughput to be able to chronologically perform two-photon calcium imaging of ALM activity over long-term (up to 6 months). By combining these approaches, we reveal stable neural representation of learned directional licking across a significant portion of a mouse’s lifetime. We also show that new task learning creates new preparatory states (Fig. 3), consistent with the motor cortex acquiring new activity patterns ^84^. In addition, we show that multiple preparatory states can encode the same action in parallel representations across distinct contexts (Fig. 5). The formation of parallel new representations protects the previously learned representations from erasure during new motor learning (Fig. 4), thus enabling stable retention of existing motor memories (Extended Data Fig. 10d). These findings significantly extend previous findings and reveal the underlying neural code for stable motor skill retention in the face of continual learning.

A combinatorial code requires high-capacity storage for motor memories due to the potentially many combinations of actions and contexts. Notably, our network modeling suggests that stable motor memory is rooted in high-dimensional representations and requires a network architecture that can readily acquire and store new activity patterns (Fig. 6e-h). It remains to be determined how such high-dimensional representations map onto neural circuits. Preparatory activity is maintained by recurrent loops between ALM and subcortical regions ^13,15,85^, including thalamus ^86–88^, midbrain ^89^, and cerebellum ^90–92^. Where might motor memories be stored? We hypothesize that the cerebellum may be a potential candidate. Cerebellar granule cells integrate inputs from the neocortex and form the basis for cerebellar output that influences preparatory activity ^14,90,91^. Cerebellar granule cells are the most numerous cell type in the brain, which could provide a substrate for high-dimensional representations with minimal interference between motor memories ^93,94^. Future work probing mechanisms of memory storage in the cerebellum may be of interest.

## Acknowledgements

We thank S. Lisberger, J. Yau, D. Herzfeld, H. Inagaki, M. Economo, D. Lipshutz, D. Ji, and members of the Li Lab for comments on the manuscript and insightful discussions. This work was funded by the Pew Scholars Program, NIH NS112312, NS113110, NS131229, NS132025, McKnight Foundation, and Simons Collaboration on the Global Brain. JHK is supported by National Research Foundation of Korea (RS-2023-00238217). KD is supported by Allen Institute for Neural Dynamics. Figure 1 diagrams were created with BioRender.com.

## Author contributions

JHK and NL conceived and designed the experiments. JHK performed the experiments. JHK analyzed data. KD performed modeling. JHK, KD, and NL wrote the paper.

## Extended Data Figures and Legends

**Extended Data Figure 1.**
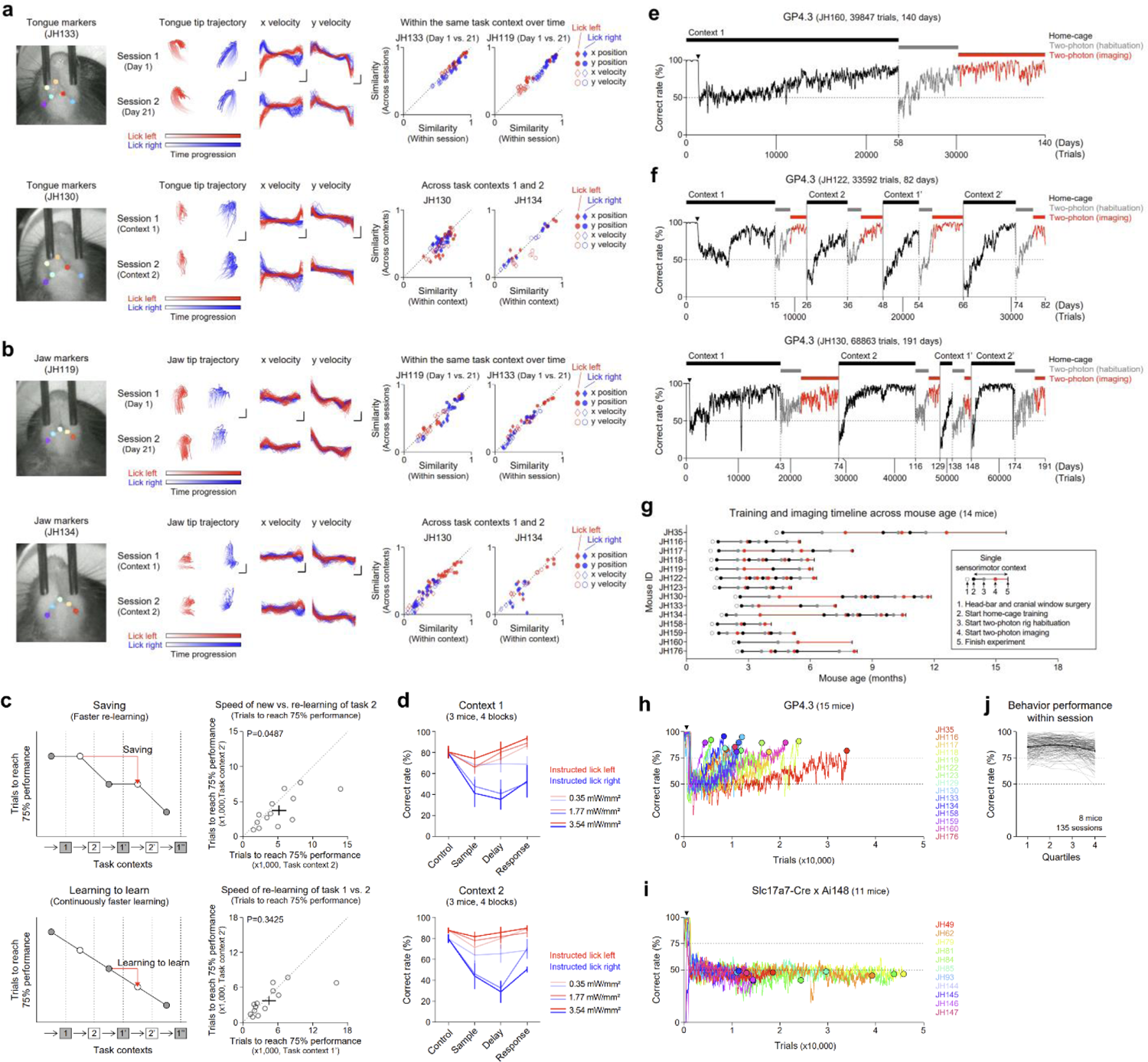
Behavioral analysis, behavioral training, and experimental timeline. **a.** Top left, representative video frame with automatically labeled tongue markers using DeepLabCut. Top middle, superimposed tongue tip trajectories and x and y velocities of individual lick events during lick left (red) and lick right (blue). Data from an example mouse across sessions within the same task context. Tongue tip trajectory scale bar, 4 pixels (x) and 6 pixels (y). X velocity scale bar, 12 ms and 2 pixels/s. Y velocity scale bar, 12 ms and 1.5 pixels/s. Top right, scatter of averaged pairwise similarity of single lick events (Pearson’s correlation) calculated within session versus across sessions. Data from two mice. Bottom, same as top but for data across task contexts 1 and 2. **b.** Same as **a**, but for jaw marker analysis. Jaw tip trajectory scale bar, 4 pixels (x) and 4 pixels (y). X velocity scale bar, 12 ms and 1 pixels/s. Y velocity scale bar, 12 ms and 1.5 pixels/s. **c.** Left, schematics of learning speed under two models. Memory-related saving effect (top): faster re-learning only for previously learned tasks. Learning-to-learn effect (bottom): faster learning each time. Right, faster reversal learning is consistent with a saving effect. Re-learning of task context 2’ is significantly faster than initial learning of task context 2 (top). P=0.0487, paired t-test. Circles indicate individual mice (N=13 mice). We examine task context 2 because the initial learning of task context 1 is confounded by the exposure to home-cage training. To examine learning-to-learn effect, we compare the speed of re-learning task context 1’ versus re-learning task context 2’ (bottom). The two conditions have similar task-specific prior training. No significant difference is observed. P=0.3425, paired t-test. **d.** Same as Fig. 1i, but separately plotting photoinhibition results for task context 1 (left) and task context 2 (right). **e.** Experimental timeline of an example mouse imaged within the same task context over extended time. Black, behavior training in automated home-cage. Gray, habituation in two-photon setup. Red, calcium imaging in two-photon setup. All the trials are concatenated. Black triangle indicates the end of learning voluntary head-fixation and start of learning in tactile instructed licking task. Averaging window, 100 trials. **f.** Same as **e**, but for two mice imaged across different task contexts. **g.** Summary plot of experimental timeline from all GP4.3 mice used for imaging in this study. **h-i**. Behavior performance curves for the initial learning from GP4.3 mice (**h**, n=15 mice, all were trained in automated home-cage) and Slc17a7-Cre x Ai148 mice (**i**, n=11 mice, 7 mice were trained in automated home-cage and 4 mice were manually trained). Different colors represent individual mice. Circles indicate end of the learning curves for GP4.3 mice and termination of training for Slc17a7-Cre x Ai148 mice. j. Behavior performance within imaging sessions across 4 segments of trials. Thin gray lines indicate individual sessions. Thick black lines indicate mean ± s.e.m. Data from Fig. 2b.

**Extended Data Figure 2.**
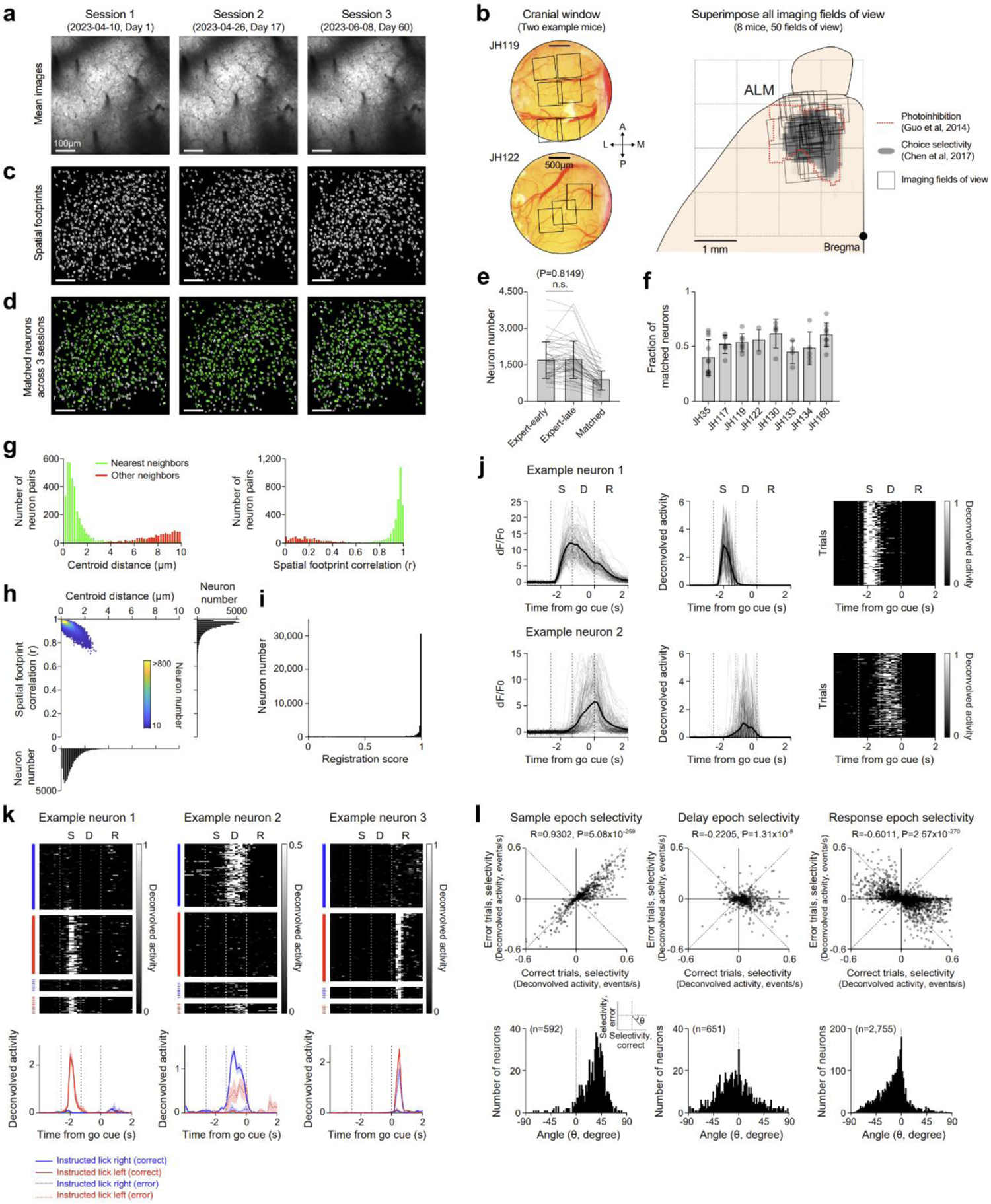
Preprocessing of imaging data and ALM preparatory activity. a. Mean two-photon fluorescence images from the same field of view (FOV) across 3 imaging sessions (Day 1, 17, and 60). b. Left, cranial windows from two example mice. Each black box indicates one imaging FOV (600 x 600 µm). Right, all imaging FOVs (n=50 from 8 mice). Imaging FOVs cover ALM, defined as the area where photoinhibition during the delay epoch impairs behavior performance (dotted red line ^61^) and exhibiting enriched choice selectivity (gray ^95^). c. Spatial footprints of individual neurons from the same FOV across 3 imaging sessions, which are the output of Suite2p (Methods). d. Identified co-registered neurons (green) across 3 imaging sessions, which are computed by CellReg (Methods). e. The number of neurons from the expert-early session (n=1,690 ± 758, mean ± SD), expert-late session (n=1,704 ± 777), and matched neurons in both expert-early and expert-late sessions (n=855 ± 402). 12.80 ± 8.90 (mean ± SD) days between imaging sessions. Data from Fig. 2. f. Fraction of match neurons across individual mice. Dots, individual FOVs. Error bars, mean ± SD. g. Distribution of centroid distance (left) and spatial footprint correlation (right) from nearest neighboring neuronal pairs (green) and other neighboring neuronal pairs within 10 µm (red). Centroid distance and spatial footprint correlation are parameters used to define co-registered neurons across imaging sessions used by CellReg package. Data from the same FOV in **a**, **c, d**. h. Density map between centroid distance and spatial footprint correlation from all co-registered neurons (n=42,739 from 8 mice). Data from Fig. 2. i. Distribution of registration score from all co-registered neurons (n=42,739 from 8 mice). j. dF/F_0_ activity (left), deconvolved activity (middle), and heatmap of single trial deconvolved activity (right) from two example neurons. Thick lines represent the mean; thin lines represent single trials. k. Single trial deconvolved activity (top) and peristimulus time histograms (PSTH, bottom) for correct and error trials are shown for three example ALM neurons. Trial types are based on instructed lick direction (blue, lick right; red, lick left). Correct trials, solid lines. Error trials, dotted lines. mean ± s.e.m. l. Top, comparison of individual neuron trial-type selectivity between correct and error trials. Neurons with significant trial-type selectivity (P<0.001, two-tailed t-test). Selectivity is the difference in deconvolved activity between instructed lick right and lick left trials during the early sample epoch (left), late delay epoch (middle), and response epoch (right). On error trials, when mice licked in the opposite direction to the instruction provided by object location (Fig. 2a), a majority of ALM neurons switched their trial type preference to predict the licking direction during the delay and response epochs, as indicated by the negative correlations (R, Pearson’s correlation). Bottom, histogram of selectivity angle between correct and error trials. A negative angle indicates neuron switching selectivity on error trials. Bin size: 2°.

**Extended Data Figure 3.**
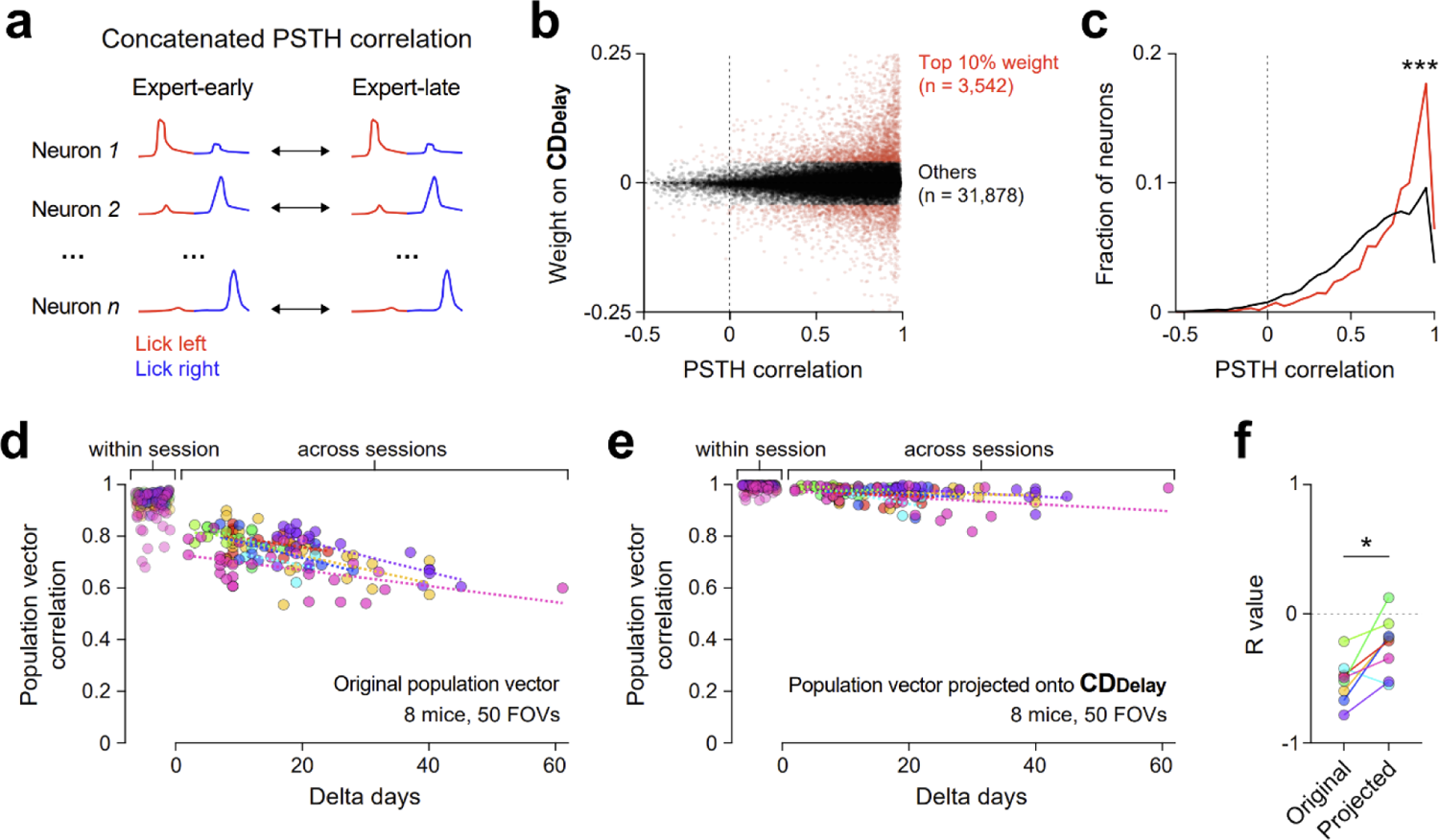
Activity drift in individual neurons and population activity. a. Quantification of PSTH stability. Pearson’s correlation between PSTHs of individual neurons from different imaging sessions (expert-early and expert-late). b. Relationship between PSTH stability and weight contribution to the **CD_Delay_**. Dots, individual neurons. Red neurons (n=3,542) are the top 10% weight contributors to the **CD_Delay_**. Black neurons (n=31,878) are the remaining 90% of neurons. c. Probability density functions of PSTH correlation for the top 10% weight contributors (red) and the other neurons (black). P=2.85×10^-114^, two-sample Kolmogorov-Smirnov test. d. Pearson’s correlation between vectors of concatenated PSTHs across the whole population as a function of delta days between expert-early and expert-late imaging sessions. Different colors represent different mice. Dotted lines indicated linear regressions of individual mice across days. e. Same as **d**, but for population activity vectors projected onto the **CD_Delay_**. f. R values of linear regressions of individual mice in panels **d** and **e**. P= 0.0156, paired t-test. Data from Fig. 2 (50 fields of view from 8 mice); mean ± s.e.m.

**Extended Data Figure 4.**
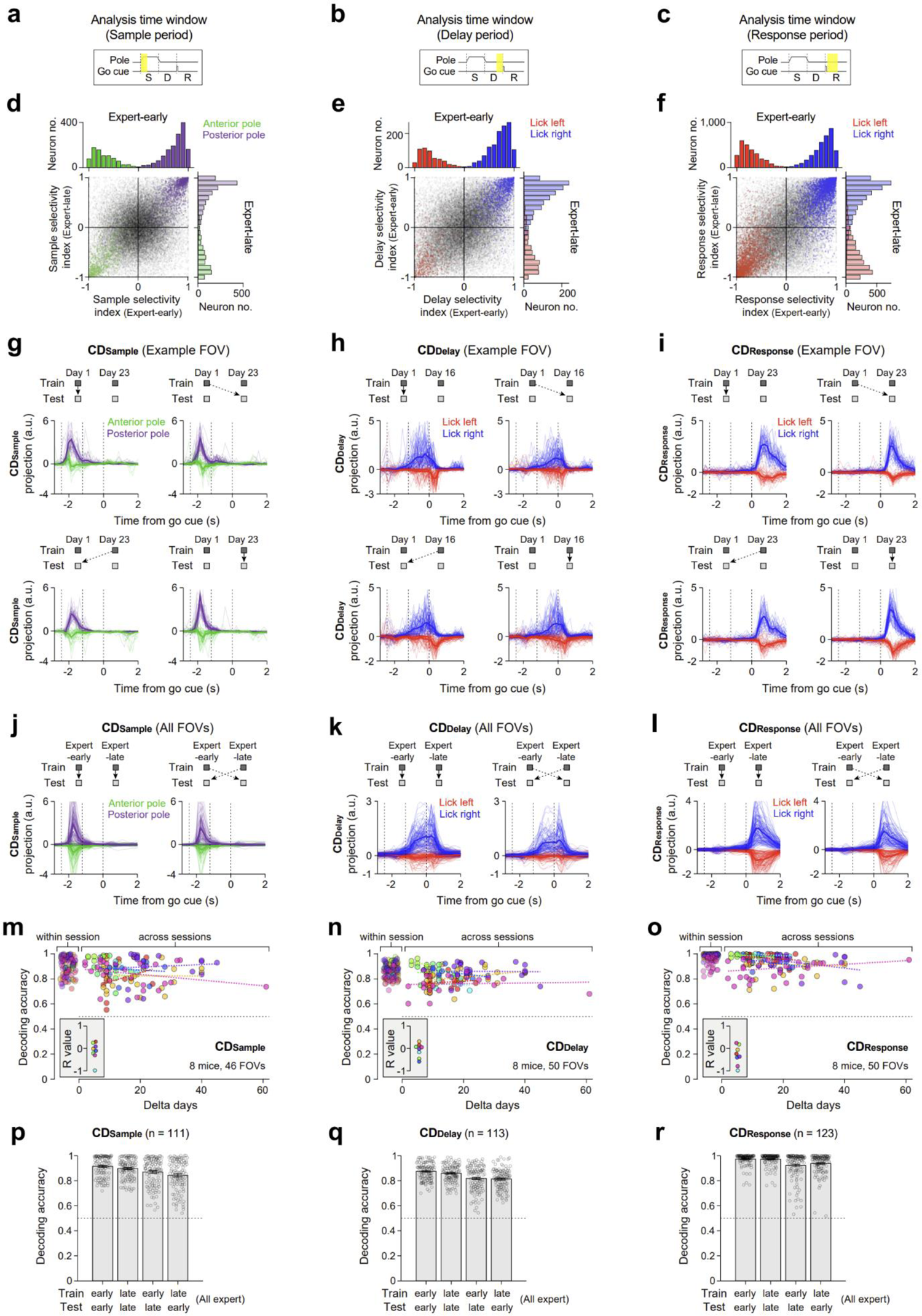
Task-related activity during the sample, delay, and response epochs within the same task context. **a-c**. Analysis time window to estimate coding direction in different task epochs. Early sample epoch (**a**, **CD_Sample_**), late delay epoch (**b**, **CD_Delay_**), and early response epoch (**c**, **CD_Response_**) were used, respectively. **d-f**. Scatter plots and histograms of individual neuron selectivity index during the sample (**d**), delay (**e**), and response epoch (**f**) comparing expert-early and expert-late sessions. Colors indicate neurons with significant trial-type selectivity (P<0.001, two-tailed t-test) during specific epochs in expert-early sessions. Neurons are colored based on their preferred trial-type in expert-early sessions. Green, neurons preferring anterior pole position. Purple, neurons preferring posterior pole position. Red, neurons preferring lick left. Blue, neurons preferring lick right. Gray, no preference neurons. Pearson’s correlation, sample epoch, R=0.9404, P=0 (**d**); delay epoch, R=0.8861, P=0 (**e**); response epoch, R=0.9001, P=0 (**f**). g. Same as Fig. 2h, but from the example FOV projected on the **CDSample** trained on day 1 (top) or day 23 (bottom) and tested on day 1 (left) or day 23 (right). h. Same as Fig. 2h (for **CD_Delay_**) replotted here for comparison. i. Same as Fig. 2h, but for **CD_Response_**. **j-l**. Trial-averaged ALM activities projected on the **CDSample** (**j**), **CD_Delay_** (**k**), and **CD_Response_** (**l**) from the same session (left) and across different sessions (right). Thin lines represent individual sessions. Thick lines represent the mean. **m-o**. Same as Fig. 2i, but for **CDSample** (**m**), **CD_Delay_** (**n**), and **CD_Response_** (**o**). P=0.4203 (**o**), P=0.4870 (**n**), P=0.0886 (**o**), R values of linear regression, t-test against 0. **p-r**. Same as Fig. 2j, but for **CDSample** (**p**, n=111 pairs of sessions, 8 mice), **CD_Delay_** (**q**, n=113 pairs of sessions, 8 mice), and **CD_Response_** (**r**, n=123 pairs of sessions, 8 mice).

**Extended Data Figure 5.**
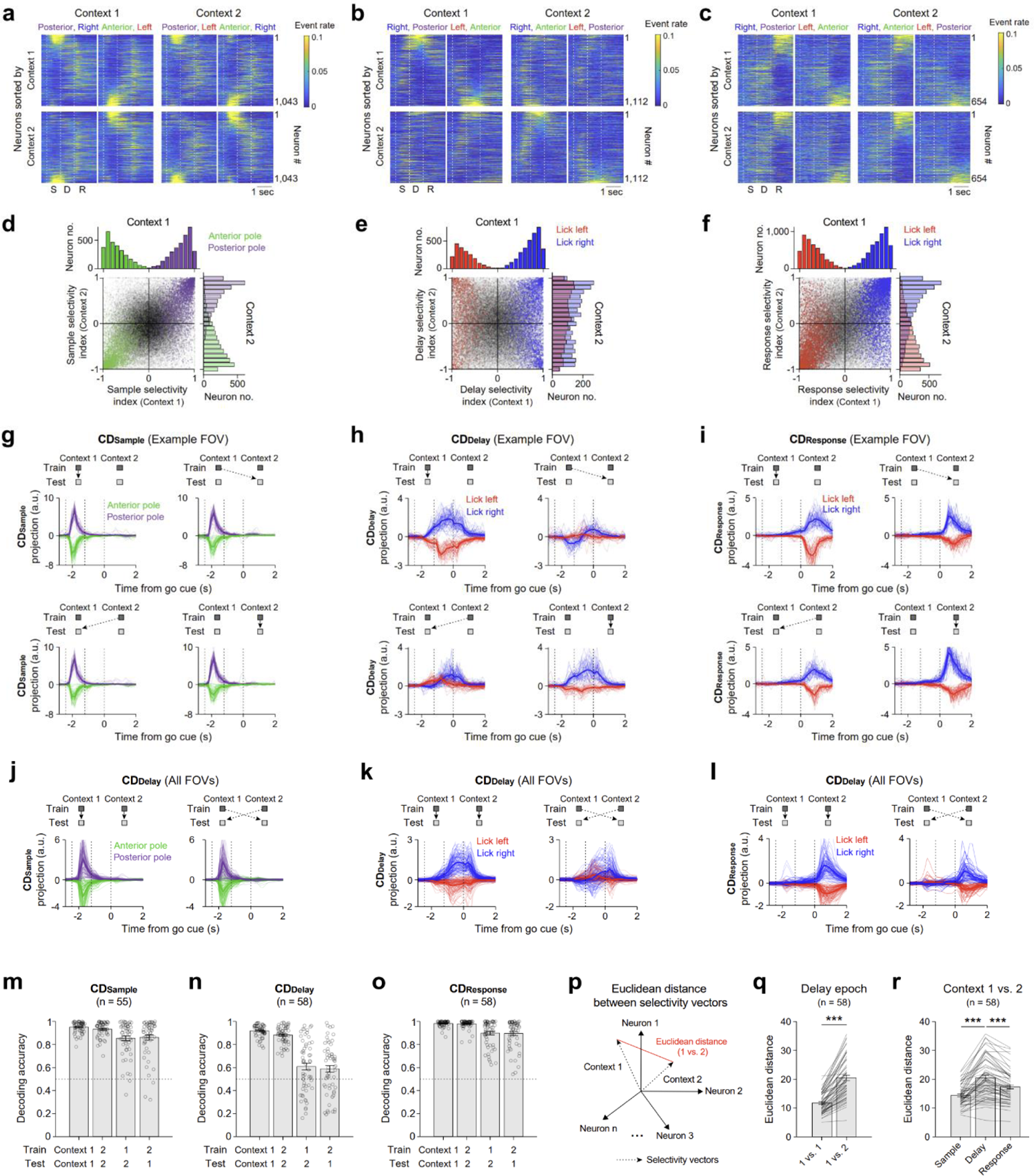
Task-related activity during the sample, delay, and response epochs across different tasks contexts. **a-c**. Same as Fig. 3c, but sorting the same neuronal population based on their selectivity during the sample epoch (**a**, n=1,043), delay epoch (**b**, n=1,112), and response epoch (**c**, n=654) in task context 1 (top) or task context 2 (bottom). **a**, **b**, and **c** contain different fields of views. **d-f**. Scatter plots and histograms of individual neuron selectivity index during the sample (**d**), delay (**e**), and response epoch (**f**) comparing task contexts 1 and 2. Colors indicate neurons with significant trial-type selectivity (P<0.001, two-tailed t-test) during specific epochs in task context 1. Neurons are colored based on their preferred trial-type in task context 1. Green, neurons preferring anterior pole position. Purple, neurons preferring posterior pole position. Red, neurons preferring lick left. Blue, neurons preferring lick right. Gray, no preference neurons. Pearson’s correlation, sample epoch, R=0.8707, P=0 (**d**); delay epoch, R=-0.0057, P=0.6774 (**e**); response epoch R=0.6804, P=0 (**f**). **g-i**. Same as Fig. 3f, but for **CDSample** (**g**), **CD_Delay_** (**h**), and **CD_Response_** (**i**). **j-l**. Trial-averaged ALM activities projected on the **CDSample** (**j**), **CD_Delay_** (**k**), and **CD_Response_** (**l**) within the same task context (left) and across different task contexts. **m-o**. Same as Fig. 3g, but for **CDSample** (**m**, n=55 pairs of sessions, 10 mice), **CD_Delay_** (**n**, n=58 pairs of sessions, 10 mice), and **CD_Response_** (**o**, n=58 pairs of sessions, 10 mice). **p.** Schematic of calculating Euclidean distance between selectivity index vectors. **q.** Euclidean distance between the delay epoch selectivity vectors calculated within task context (1 vs. 1) and across task contexts (1 vs. 2). Selectivity vectors within task context are calculated using split-half trials from the same session. ***P=4.03×10^-20^, paired t-test. Mean ± s.e.m. **r.** Euclidean distance of sample, delay, and response epoch selectivity vectors across task contexts (1 vs. 2). Sample vs. delay epoch, ***P=9.21×10^-13^; delay vs. response epoch, ***P=1.71×10^-12^, paired t-test. Mean ± s.e.m.

**Extended Data Figure 6.**
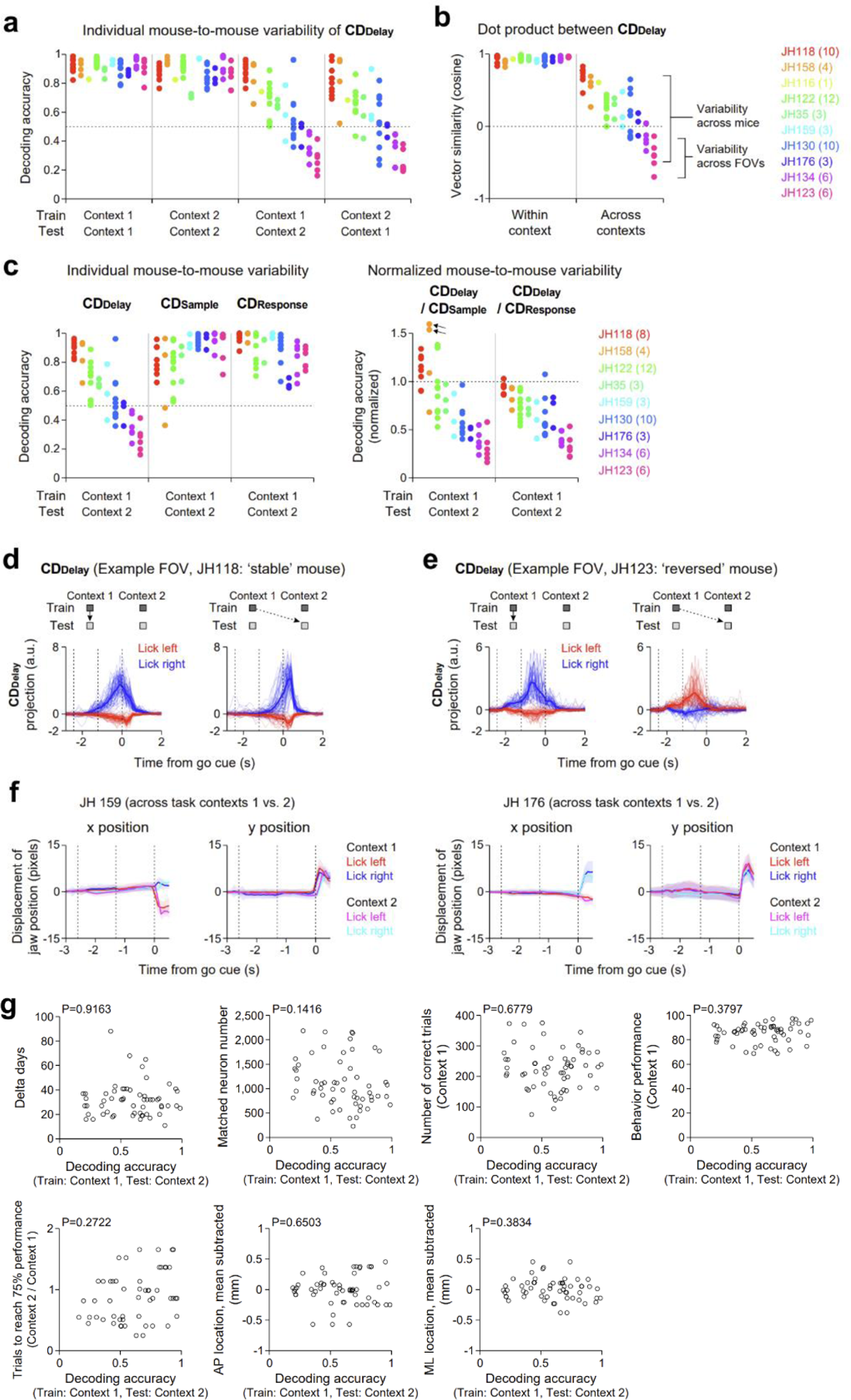
Individual variability across mice in the degree of CD_Delay_ reorganization across task contexts. a. Data from Fig. 3g, but broken out by individual mice, sorted by mean decoding accuracy across task contexts (train context 1 and test context 2). Individual mice are plotted separately in different colors. b. Dot product between **CD_Delay_** within the same task context (left) and across different task contexts (right). Individual mice are sorted by mean dot product of **CD_Delay_** across task contexts. Same color scheme as **a**. Note that variability across mice is much higher than variability across FOVs within the same mouse. Numbers in brackets indicate pairs of sessions for each mouse. c. Left, decoding accuracy of the **CD_Delay_**, **CD_Sample_**, and **CD_Response_** across task contexts. Right, decoding accuracy of the **CD_Delay_**, normalized by that of **CD_Sample_** and **CD_Response_**. Arrows indicate two outlier data points (y-axis values, 1.93 and 2.23). **d-e**. Same as Fig. 3f (top), but data from an example FOV of a mouse with stable **CD_Delay_** across task contexts (**c**, JH118; see **a-b**) and a mouse with reversed **CD_Delay_** (**d**, JH123; see **a-b**). f. Displacement of x and y jaw positions in task contexts 1 and 2 from two example mice. Mean ± SD across trials. g. No relationship between decoding accuracy of **CD_Delay_** across task contexts (x axis) versus 7 parameters (y axis) as follows: delta days between imaging sessions; matched number of neurons co-registered across imaging sessions; number of correct trials from task context 1; behavior performance from task context 1; relative learning speed to reach 75% behavior performance (number of trials to reach criterion performance in task context 2 relative to task context 1); relative AP location of imaging FOVs after subtracting mean AP location of each mouse; relative ML location of imaging FOVs after subtracting mean ML location of each mouse.

**Extended Data Figure 7.**
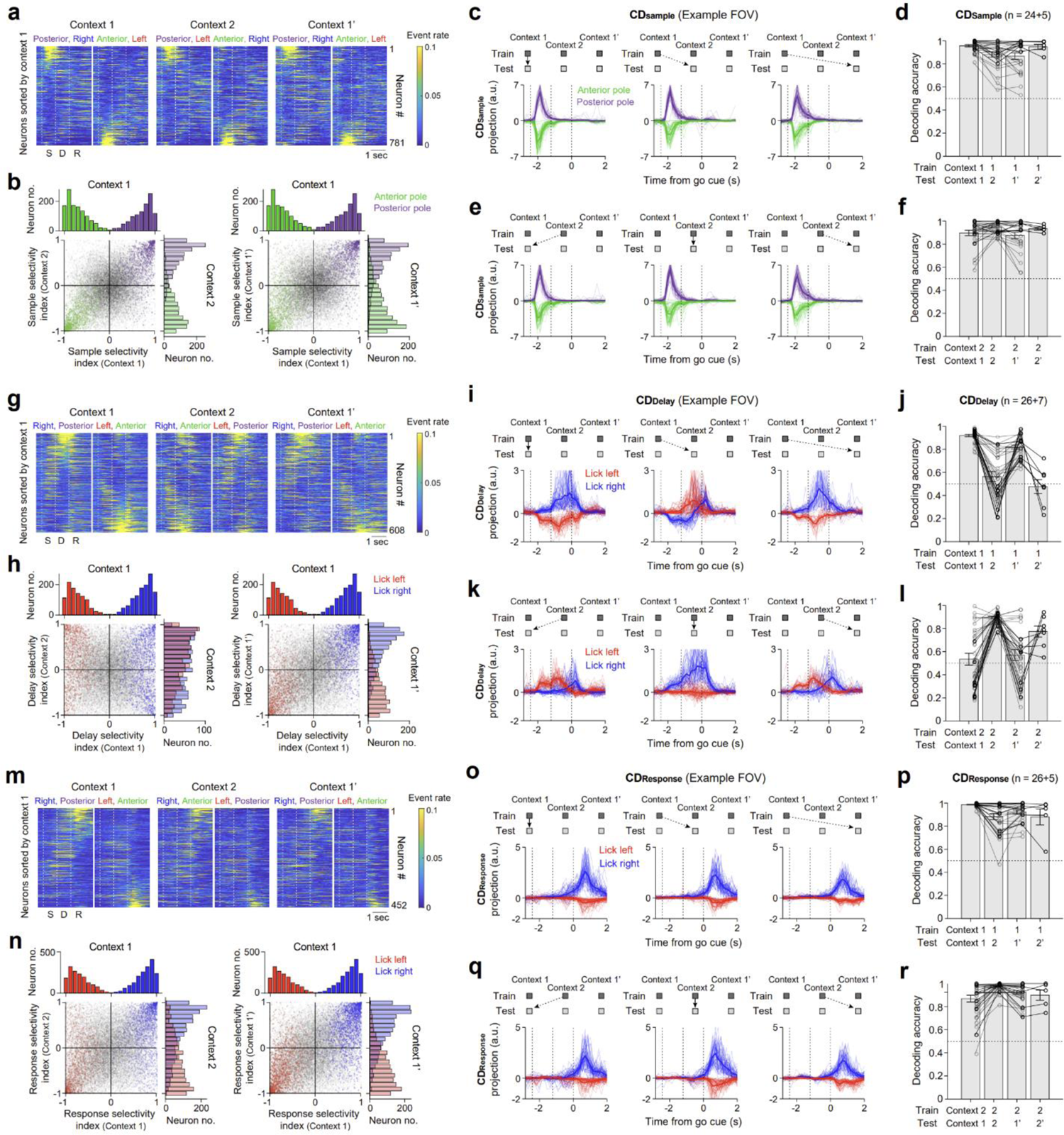
Task-related activity during the sample, delay, and response epochs across task context 1, 2, and 1’. **a.** Mean deconvolved activities from an example field of view across three task contexts (n=781 neurons). **b.** Scatter plots and histograms of individual neuron selectivity index during the sample epoch comparing task contexts 1 and 2 (left) or task contexts 1 and 1’ (right). Colors indicate neurons with significant trial-type selectivity (P<0.001, two-tailed t-test) during specific epochs in expert-early sessions. Neurons are colored based on their preferred trial-type in task context 1. Green, neurons preferring anterior pole position. Purple, neurons preferring posterior pole position. Gray, no preference neurons. Pearson’s correlation, task context 1 vs. 2, R=0.8912, P=0; task context 1 vs. 1’, R=0.8290, P=0. **c-f.** Same as Fig. 4f-i, but for activity during the sample epoch. In **d** and **f**, gray circles and lines indicate FOVs imaged across task contexts 1, 2, and 1’ (n=24 FOVs, 5 mice); black circles and lines indicate subset of FOVs imaged across task contexts 1, 2, 1’, and 2’ (n=5 FOVs, 2 mice). **g-l**. Same as **a**-**f**, but for activity during the delay epoch. **h**, red indicates neurons preferring lick left and blue indicates neurons preferring lick right in task context 1. Pearson’s correlation, task context 1 vs. 2, R=-0.1224, P=1.38×10^-8^; task context 1 vs. 1’, R=0.7675, P=0. In **j** and **l**, FOVs imaged across task contexts 1, 2, and 1’ (n=26 FOVs, 5 mice); FOVs imaged across task contexts 1, 2, 1’, and 2’ (n=7 FOVs, 3 mice). Data from Fig. 4, replotted here for comparison. **m-r**. Same as **g**-**l**, but for activity during the response epoch. **n**, Pearson’s correlation, task context 1 vs. 2, R=0.6220, P=0; task context 1 vs. 1’, R=0.7624, P=0. In **p** and **r**, FOVs imaged across task contexts 1, 2, and 1’ (n=26 FOVs, 5 mice); FOVs imaged across task contexts 1, 2, 1’, and 2’ (n=5 FOVs, 3 mice).

**Extended Data Figure 8.**
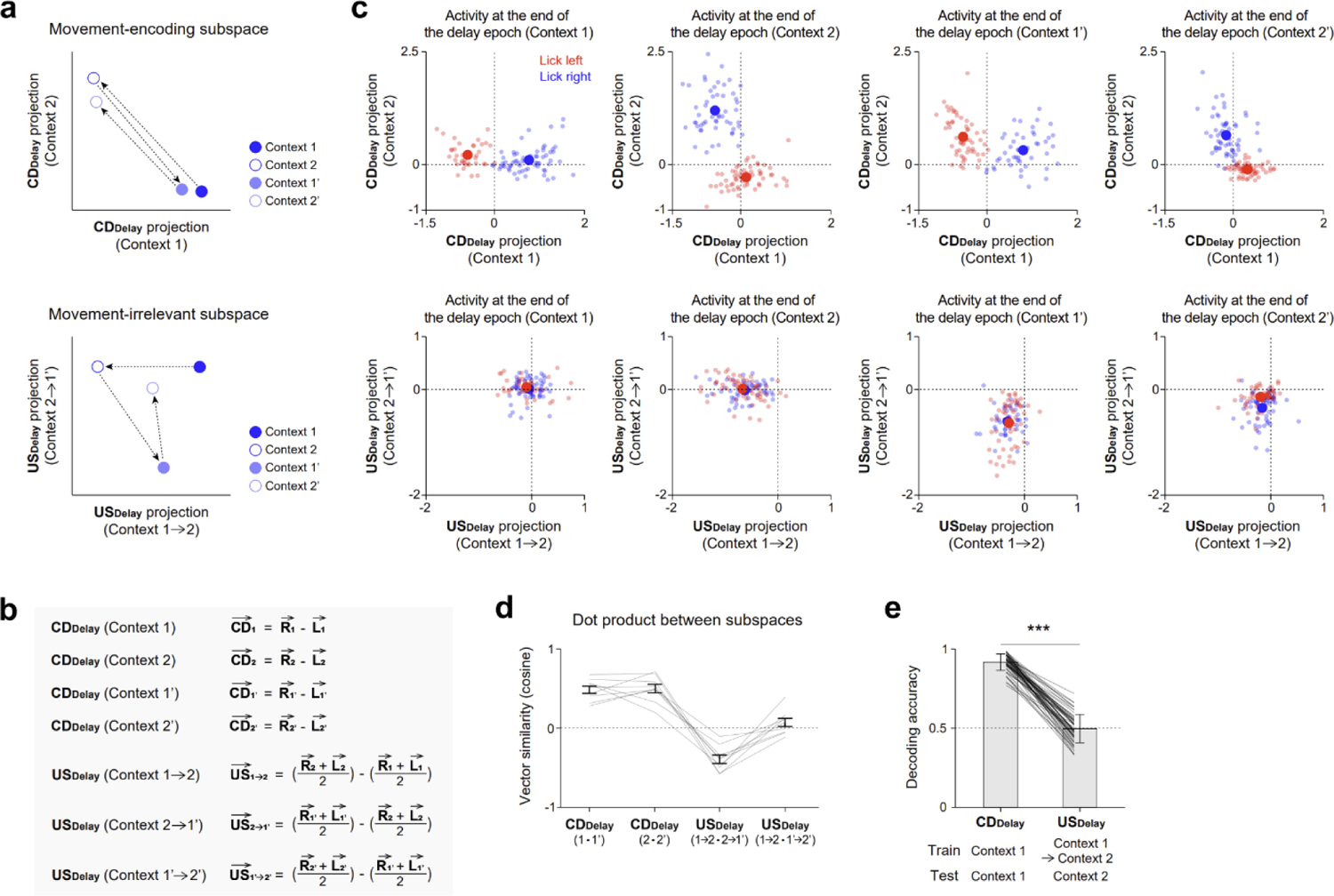
Neural activity change in movement-irrelevant activity subspace across task contexts. a. Schematic of activity changes across task contexts along coding directions (top, **CD_Delay_**, estimated from task contexts 1 and 2) and movement-irrelevant subspace (bottom, **US_Delay_**, estimated from task context 1→2 and 2→1’). b. Formula to calculate the **CD_Delay_**’s and the **US_Delay_**’s. **CD_Delay_**’s are calculated separately for each task context. **US_Delay_**’s are calculated separately for each task context change. c. Activity of an example FOV across 4 task contexts (1, 2, 1’, and 2’). Top, activity projections on the **CD_Delay_**’s. The **CD_Delay_** of task context 2 is orthogonalized to the **CD_Delay_** of task context 1 here for visualization purposes. Similar patterns of activity are re-activated in the same task context (1 vs. 1’ and 2 vs. 2’). Big solid circles represent the mean; small transparent circles represent activity in single trials. Bottom, activity projections on the **US_Delay_**’s. In contrast to the activity along the **CD_Delay_**’s, activity along the **US_Delay_**’s does not show reliable re-activation in the same context. d. Dot products (mean ± s.e.m.) between the **CD_Delay_**’s and **US_Delay_**’s (n=9 fields of view, 3 mice). Activity along the coding directions shows reliable re-activation (consistent **CD_Delay_**’s for task context 1 vs. 1’ and 2 vs. 2’). Activity in the movement-irrelevant subspace does not show consistent changes across re-learning of previous task contexts (1→2 vs. 2→1’ and 1→2 vs. 1’→2’). e. Decoding accuracy of the **CD_Delay_** and **US_Delay_** to predict lick directions. ***P=5.64×10^-39^, paired t-test. Activity projection on the **US_Delay_** does not predict lick direction.

**Extended Data Figure 9.**
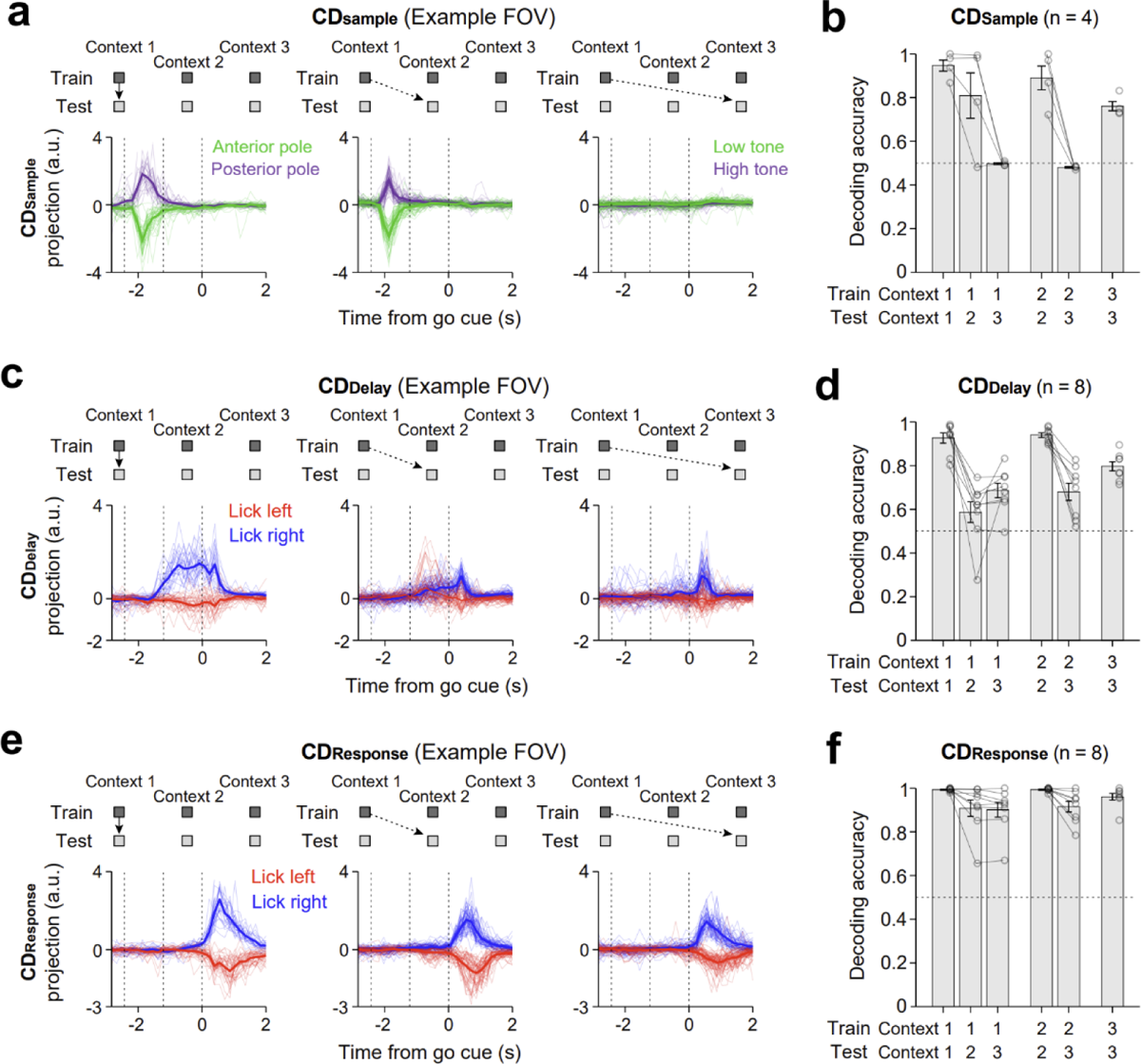
Task-related activity during the sample, delay, and response epochs across task context 1, 2, and 3. **a-b**. Same as Fig. 5e-f, but for activity during the sample epoch (n=4 fields of view, 2 mice). **c-d**. Same as Fig. 5e-f, replotted here for comparison. **e-f**. Same as Fig. 5e-f, but for activity during the response epoch (n=8 fields of view, 3 mice).

**Extended Data Figure 10.**
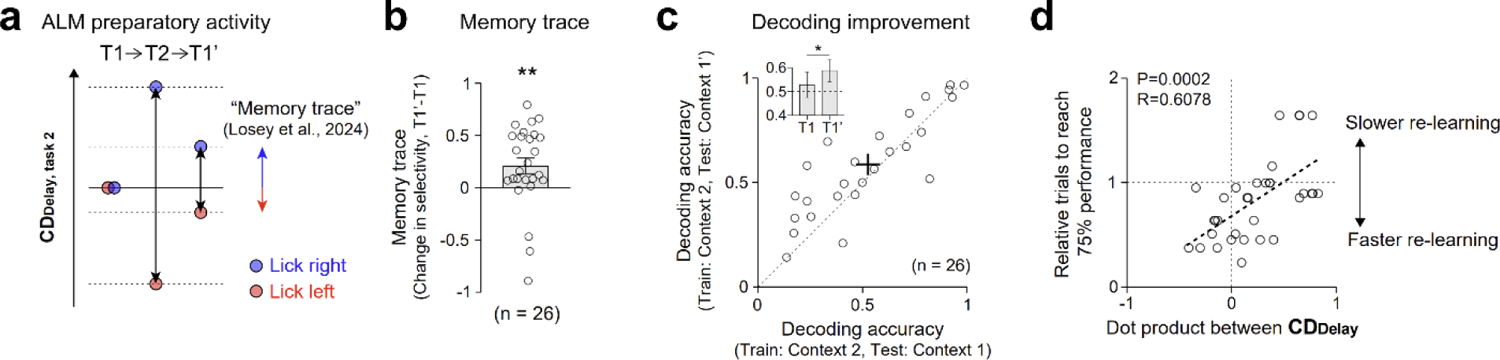
Context-specific preparatory activity retains memory trace of previous learning and reduces interference. a. Schematic of the memory trace. ALM preparatory activity from task contexts 1, 2, and 1’ is projected onto the **CD_Delay_** from task context 2. Memory trace is defined as a selectivity increase along the **CD_Delay_** for task context 2 during performance of task context 1’, as shown in black arrows combining blue and red arrows. b. Memory trace. Change in delay epoch selectivity along the **CD_Delay_** for task context 2 from task context 1 to 1’. **P=0.005, paired t-test. c. Decoding accuracy of the **CD_Delay_** for task context 2 tested on task contexts 1 (52.75 ± 5.24%) and 1’ (58.66 ± 4.63%). Cross, mean ± s.e.m. Inset, decoding accuracy in each task. *P=0.0199, paired t-test. 26 fields of view from 5 mice. d. Speed of re-learning task context 1’ as a function of the **CD_Delay_** reorganization across task contexts 1 and 2. Number of trials to reach criterion performance in task context 1’ relative to number of trials during the initial learning of task context 1. Mice exhibiting more distinct **CD_Delay_**’s across task contexts (i.e. lower dot product) re-learned the previously learned task context 1’ faster (i.e. fewer trials to reach 75% performance criterion). Each dot shows one field of view from one mouse. Dotted line, linear regression; R, Pearson’s correlation.

**Extended Data Figure 11.**
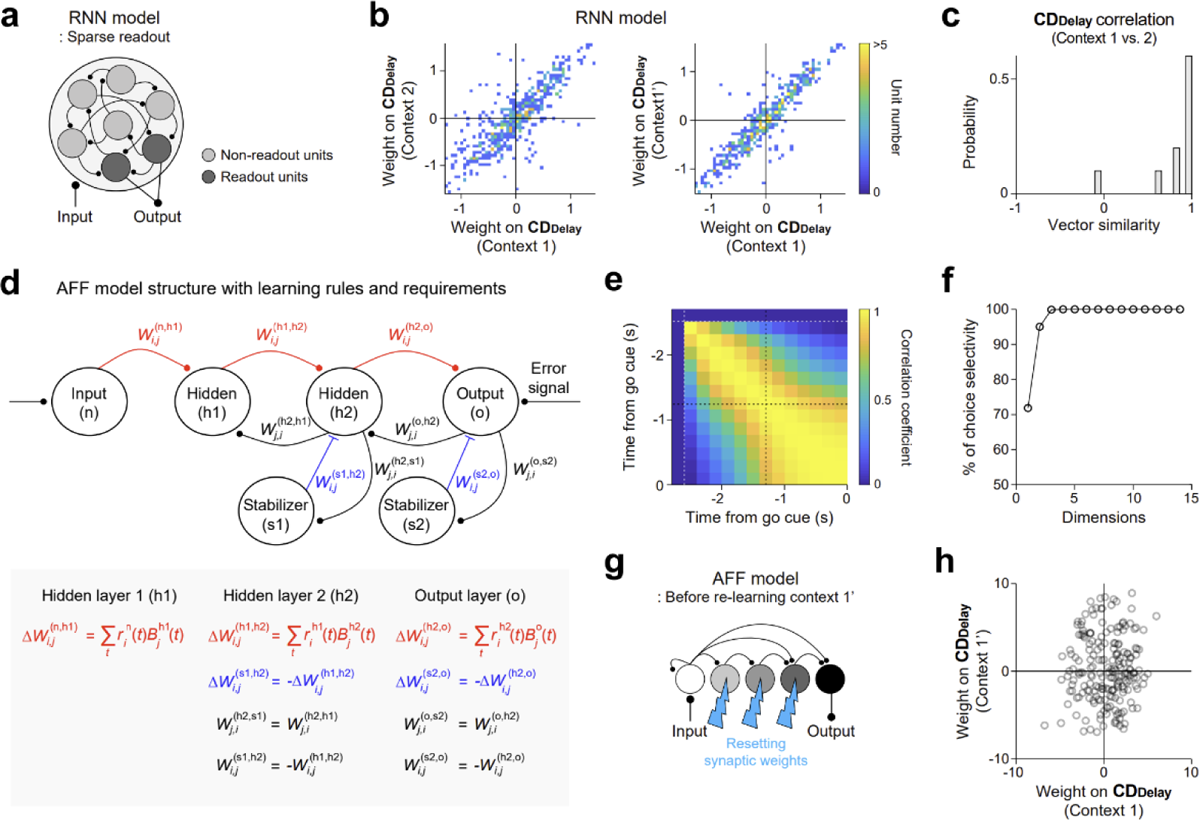
Recurrent neural network (RNN) and amplifying feedforward (AFF) network models. **a.** A schematic of RNN model with sparse readout. Only two internal units directly contribute to the output. **b-c**. Same as Fig. 6c-d, but for RNNs with sparse readout. **d.** A schematic of the AFF networks and governing equations. **e.** Analysis of the AFF network similar to Daie et al^77^. We identified directions in activity space at different time points that influence network activity along the **CD_Delay_** at the end of the delay epoch (t = 0 s). We refer to these as transitional directions. The plot shows correlation between transitional directions at time point t vs time point t’ for all sample and delay epoch time points. AFF networks generate persistent activity by passing activity through a chain of network states, where early layers influence activity in the late layers. This results in network activity sequentially traversing multiple directions in activity space, as indicated by the low correlation values off diagonal. **f.** Dimensionality of trial-type selectivity in the AFF networks. 3 dimensions captured most of the network selectivity. **g.** Resetting AFF network hidden layer weights to random values before re-learning task context 1’. **h.** Resetting synaptic weights before re-learning task context 1’ prevented the re-activation of the **CD_Delay_**. Weight contribution of the AFF units to the **CD_Delay_**’s from task contexts 1 and 1’.

**Extended Data Figure 12.**
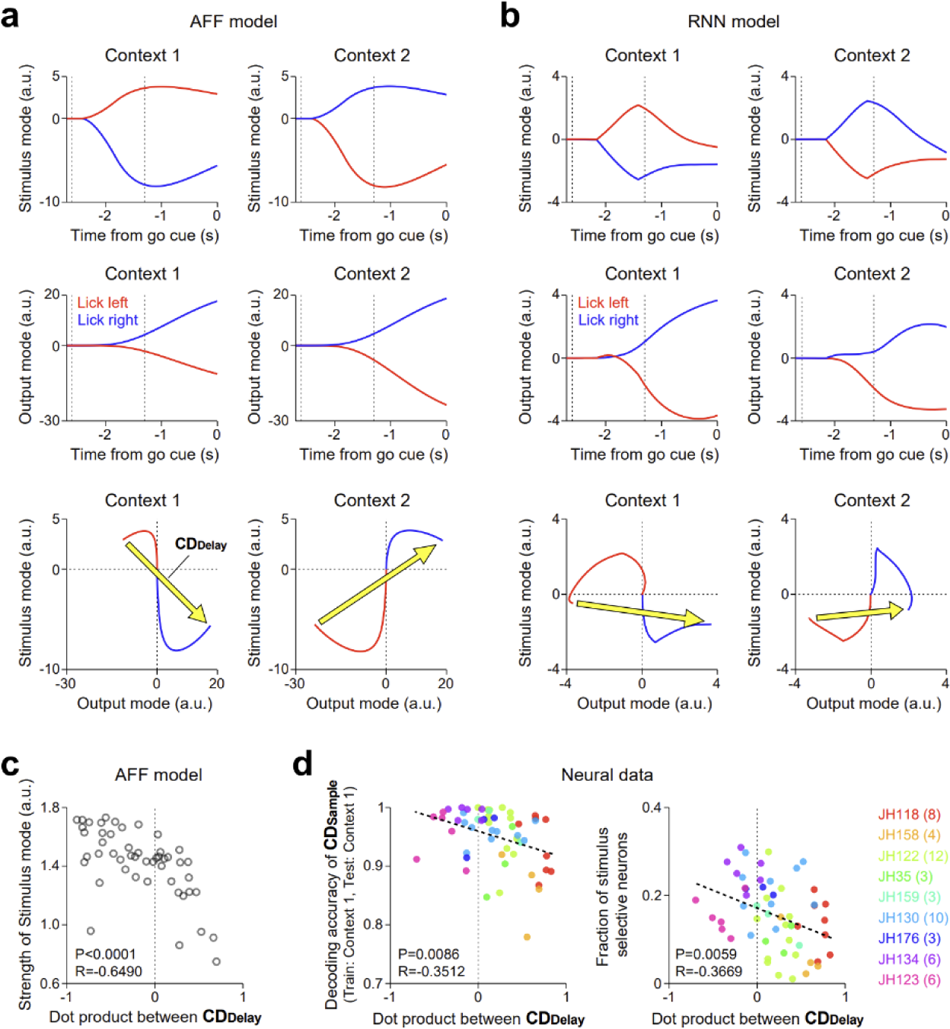
Neural dynamics within RNN and AFF networks. a. AFF network activity projected on the stimulus mode (top), output mode (middle), and in state space (bottom). Trial types are defined by lick direction. Blue, lick right. Red, lick left. See Methods for decomposition of network activity modes. AFF networks exhibit persistent activity along the stimulus mode, which combines with the ramping output mode to produce distinct **CD_Delay_**’s in each task context (yellow arrows). b. Same as **a**, but for RNN. RNNs do not maintain persistent activity along the stimulus mode, which results in stable **CD_Delay_**’s across task contexts that are aligned to the output mode. c. The strength of network activity along the stimulus mode predicts the degree of **CD_Delay_** reorganization across task contexts (dot product of the **CD_Delay_**’s from task contexts 1 and 2). Dots, individual randomly initialized AFFs. d. Neural data. Individual variability of **CD_Delay_** reorganization across task contexts is predicted by the strength of stimulus activity in ALM. The strength of ALM stimulus activity is quantified as the decoding accuracy of the **CD_Sample_** in task context 1 (trained and tested within task context), or the fraction of neurons with significant trial-type selectivity during the sample epoch in task context 1. Dots, individual FOVs. Individual mice are plotted separately in different colors.

## Methods

### Mice

This study was based on data from 36 mice (age > postnatal day 60, both male and female mice). 15 GP4.3 mice (Thy1-GCaMP6s; Jackson laboratory, JAX 024275) were used for longitudinal two-photon calcium imaging. Among them, one mouse was removed from subsequent neuronal data analyses due to the low number of matched neurons across days (see *Preprocessing of two-photon imaging data* below). 5 GAD2-ires-cre mice (JAX 010802) were used for ALM photoinhibition in home-cage. 5 additional GAD2-ires-cre mice were used only for behavior training in home-cage. 11 Slc17a7-cre mice (JAX 023527), crossed to cre-dependent GCaMP6f reporter Ai148 mice (JAX 030328) were used for behavior training but were not used for calcium imaging due to poor behavioral performance (Extended Data Fig. 1i).

All procedures were in accordance with protocols approved by the Institutional Animal Care and Use Committees at Baylor College of Medicine. Mice were housed in a 12:12 reversed light/dark cycle and tested during the dark phase. On days not tested, mice received 0.5–1 ml of water. On other days, mice were tested in experimental sessions lasting 1–2 h where they received all their water (0.5–1 ml). If mice did not maintain a stable body weight, they received supplementary water ^96^. All surgical procedures were carried out aseptically under 1–2% isoflurane anesthesia. Buprenorphine Sustained Release (1 mg/kg) and Meloxicam Sustained Release (4 mg/kg) were used for preoperative and postoperative analgesia. A mixture of bupivacaine and lidocaine was administered topically before scalp removal. After surgery, mice were allowed to recover for at least 3 days with free access to water before water restriction.

### Surgery

Mice were prepared with a clear-skull cap and a headpost ^61,96^. The scalp and periosteum over the dorsal skull were removed. For ALM photoinhibition in GAD2-ires-cre mice, AAV8-Ef1a-DIO-ChRmine-mScarlet^70^ (Stanford Gene Vector and Virus Core; titer 8.44×10^12^ vg/mL) was injected in the left ALM (anterior 2.5 mm from bregma, lateral 1.5 mm, depth 0.5 and 0.8 mm, 200 nL at each depth) using a Nanoliter 2010 injector (World Precision Instruments) with glass pipettes (20–30 µm diameter tip and beveled). A layer of cyanoacrylate adhesive was applied to the skull. A custom headpost was placed on the skull and cemented in place with clear dental acrylic. A thin layer of clear dental acrylic was applied over the cyanoacrylate adhesive covering the entire exposed skull.

For two-photon calcium imaging in GP4.3 mice, a glass window was additionally implanted over ALM. A circular craniotomy with diameter 3.2 mm was made over the left ALM (anterior 2.5 mm from bregma, lateral 1.5 mm). Dura inside craniotomy was removed. A glass assembly consisting of a single 4 mm diameter coverslip (Warner Instruments; CS-4R) on the top of two 3 mm diameter coverslips (Warner Instruments; CS-3R) was combined using optical adhesive (Norland Products; NOA 61) and UV light (Kinetic instruments Inc.; SpotCure-B6). The glass window was affixed to the surrounding skull of craniotomy using cyanoacrylate adhesive (Elmer; Krazy glue) and dental acrylic (Lang Dental Jet Repair Acrylic; 1223-clear).

### Behavior tasks and training in home-cage

Details of behavior task and training in the autonomous home-cage system have been described previously ^65^. In brief, a headport (∼20×20 mm) was in the frontal side of the home-cage. The two sides of the headport were fitted with widened tracks that guided a custom headpost (26.5 mm long, 3.2 mm wide) into a narrow spacing where the headpost could trigger two snap action switches (D429-R1ML-G2, Mouser) mounted on both sides of the headport. Upon switch trigger, two air pistons (McMaster; 6604K11) were pneumatically driven (Festo; 557773) to clamp the headpost. A custom 3D-printed platform was placed inside the home-cage in front of the headport. The stage was embedded with a load cell (Phidgets; CZL639HD) to record mouse body weight. This body weight-sensing stage was also used to detect struggles during head-fixations and triggered self-release. A lickport with two lickspouts (5 mm apart) was placed in front of the headport. Each of the lickspout was electrically coupled to the custom circuit board that detected licks via completion of an electrical circuit upon licking contacts ^61,97^. Water rewards were dispensed by two solenoid valves (The Lee Company; LHDA1233215H). The sensory stimulus for the tactile-instructed licking task was a mechanical pole (1.5 mm diameter) on the right side of the headport. The pole was motorized by a linear motor (Actuonix; L12-30-50-12-I) and presented at different locations to stimulate the whiskers. The sensory stimuli for the auditory-instructed licking task were pure tones (2 kHz or 10 kHz) provided by a piezo buzzer (CUI Devices; CPE-163) placed in front of the headport. The auditory ‘go’ cue (3.5 kHz) in both tactile and auditory tasks was provided by the same piezo buzzer.

Protocols stored on microcontrollers (Arduino; A000062) operated the home-cage system and autonomously trained mice in voluntary head-fixation and behavioral tasks, as well as carrying out optogenetic testing. In brief, mice were placed inside the home-cage and could freely lick both lickspouts that were placed inside the home-cage through the headport. The rewarded lickspout alternated between the left and right lickspouts (3 times each) to encourage licking on both lickspouts. This phase of the training acclimated mice to the lickport and the lickport was gradually retracted into the headport away from the home-cage. The lickport retraction continued until the tip of the lickspouts was approximately 14 mm away from the headport. At this point, mice could only reach the lickspouts by entering the headport with the headpost triggering the head-fixation switches. After 30 successful voluntary head-fixation switch triggers, the pneumatic pistons were activated to clamp the headpost upon the switch trigger (‘voluntary head-fixation’; Fig. 1c). The head-fixation training protocol continuously increased the pneumatic clamping duration (from 3 s to 30 s). This clamping was self-released when the body weight readings from the load-sensing platform exceeded either an upper (30 g) or lower (−1 g) threshold. Overt movements of the mice during the head-fixation typically produced large fluctuations in weight readings exceeding the thresholds. These thresholds were dynamically adjusted during the training process.

When mice successfully performed head-fixation training protocol by reaching 30 s head-fixation duration, the next training protocol for the tactile-instructed licking task began. In the tactile-instructed licking task, mice used their whiskers to discriminate the location of a pole and reported choice using directional licking for a water reward (Fig. 1d) ^61,96^. The pole was presented at one of two positions that were 6 mm apart along the anterior-posterior axis. The posterior pole position was approximately 5 mm from the right whisker pad. The sample epoch was defined as the time between the pole movement onset to 0.1 s after the pole retraction onset (sample epoch, 1.3 s). A delay epoch followed during which the mice must keep the information in short-term memory (delay epoch, 1.3 s). An auditory ‘go’ cue (0.1 s duration) signaled the beginning of response epoch and mice reported choice by licking one of the two lickspouts. Task training had three subprotocols that shaped mice behavior in stages. First, a ‘directional licking’ subprotocol trained mice to lick both lickspouts and switch between the two. Then, a ‘discrimination’ subprotocol taught mice to report pole position with directional licking. Finally, a ‘delay’ subprotocol taught mice to withhold licking during the delay epoch and initiate licking upon the ‘go’ cue by gradually (in 0.2 s steps) increasing the delay epoch duration up to 1.3 s. At the end of the delay subprotocol, the head-fixation duration was further increased from 30 s to 60s. The head-fixation duration was increased by 2 s after every 20 successful head-fixations. This was done to obtain more behavioral trials in each head-fixation. The program also adjusted the probability of each trial type to correct biased licking of the mice.

Mice were first trained in one sensorimotor contingency (Fig. 1b, task context 1; anterior pole position→lick left, posterior pole position→lick right). Then, the correspondence between pole locations and lick directions was reversed (task context 2; anterior pole position→lick right, posterior pole position→lick left). Over multiple months, mice could learn multiple rounds of sensorimotor contingency reversal depending on experiment (see *Performance criteria for contingency reversals and acclimation to imaging setup* below).

For auditory-instructed licking task, mice were trained to perform directional licking to report the frequency of a pure tone presented during the sample epoch (Fig. 5b, task context 3; 2 kHz (low tone)→lick left, 10 kHz (high tone) →lick right). Task structures such as the delay epoch (1.3 s) and auditory go cue (3.5 kHz, 0.1 s) were the same as the tactile-instructed licking task.

### Performance criteria for contingency reversals and acclimation to imaging setup

For mice that underwent optogenetic experiment in home-cage, contingency reversal was automatically introduced when mice reached performance criteria of >75% correct and <50% early lick for 100 trials in a given task contingency (Figs. 1e and 1h). Mice learned multiple rounds of contingency reversals before optogenetic experiment initiated. Optogenetic experiment was manually initiated based on inspections of behavioral performance (Fig. 1h).

Mice for two-photon imaging were over-trained in each task context to reach performance criteria of >80-85% correct for 100 trials. Over-training facilitated faster habituation after transferring to the two-photon setup. After mice acquired this high level of task performance in home-cage training, we transferred the mice to the imaging setup where they performed the same task in daily sessions under the two-photon microscope. During this period, mice were singly housed outside of the automated home-cage system. A brief acclimation period lasting for a few days was required to habituate the mice to perform the task under the microscope (Extended Data Fig. 1e-g). We started imaging sessions once mice recovered their task performance (typically >75%). After imaging across multiple sessions, mice were returned to the automated home-cage again in which they learned other tasks. In this manner, we repeatedly transferred mice between the automated home-cage and two-photon setup for as long as possible (Extended Data Fig. 1f-g).

For tactile-instructed licking task, mice were first trained and imaged in one sensorimotor contingency (Fig. 3b, task context 1). After imaging under the two-photon microscope, we transferred the mice back to the home-cage and reversed the sensorimotor contingency (Fig. 3b, task context 2). The mice were over-trained in the new task contingency before transferring to the two-photon setup to re-image the same ALM populations across task contexts (task context 1→2; 10 mice). In a subset of mice, after imaging, we re-trained the mice in the previous contingency in the home-cage (Fig. 4b, task context 1’). After achieving proficient task performance, we translocated the mice to the two-photon setup and imaged the same ALM populations again (task context 1→2→1’; 5 mice). In a subset of mice, we further repeated the contingency reversal one more time and imaged across four task contexts (task context 1→2→1’→2’; 3 mice).

For auditory-instructed licking task, mice were imaged first in the tactile task contexts 1 and 2 before training in the auditory task to image the same ALM populations across task contexts (task context 1→2→3; 8 mice).

### ALM photoinhibition in home-cage

The procedure for ALM photoinhibition in home-cage has been described previously ^65^. Light from a 633 nm laser (Ultralaser; MRL-III-633L-50 mW) was delivered via an optical fiber (Thorlabs; M79L005) placed above the headport (Fig. 1g). Photostimulation of the virus injection site was through a clear skull. The photostimulus was a 40 Hz sinusoid lasting for 1.3 s, including a 100 ms linear ramp during photostimulus offset to reduce rebound neuronal activity^98^. Photostimulation was delivered in a random subset of trials (18%) during either the sample, delay, or response epoch. Photostimulation started at the beginning of the task epoch.

Photostimulation power was 2.5, 12.5, or 25 mW, randomly selected in each trial. Therefore, the probability of each photostimulation condition was 2% (total of 9 conditions). The size of the light beam on the skull surface was 7.07 mm^2^ (3.0 mm diameter). 2.5, 12.5, and 25.0 mW power corresponded to 0.35, 1.77, and 3.54 mW/mm^2^ in light intensity. This range of the light intensity was much lower than the previous studies ^61,62^ (typically 1.5 mW with a light beam diameter of 0.4 mm, corresponding to 11.9 mW/mm^2^). To prevent the mice from distinguishing photostimulation trials from control trials using visual cues, a masking flash was delivered using a 627 nm LED on all trials near the eyes of the mice. The masking flash began at the start of the sample epoch and continued through the end of the response epoch in which photostimulation could occur.

### Videography

Two CMOS cameras (Teledyne FLIR; Blackfly BFS-U3-04S2M) were used to measure orofacial movements of the mouse from the bottom and side views (Extended Data Figs. 1a-b and 5e). Both the bottom and side views were acquired at 224 x 192 pixels and 400 frames/s. Mice performed the task in complete darkness, and videos were recorded under infrared 940 nm LED illumination (Luxeon Star; SM-01-R9). A custom written software controlled the video acquisition ^99^.

### Two-photon imaging

A Thorlabs Bergamo II two-photon microscope equipped with a tunable femtosecond laser (Coherent; Chameleon Discovery) is controlled by ScanImage 2016a (Vidrio). GCaMP6s was excited at 920 nm. Images were collected with a 16X water immersion lens (Nikon, 0.8 NA, 3 mm working distance) at 2x zoom (512 x 512 pixels, 600 x 600 µm). For all imaging sessions, we performed volumetric imaging by serially scanning five planes (30 or 40 μm equally spaced along z-axis) at 6 Hz each. The range of depth from all imaging planes was 120–500 μm below the pial surface, and the range of laser power was 80-225 mW, measured below the objective. To identify the spatial locations of individual field of view (FOV), we imaged at the pial surface before imaging during the task (Extended Data Fig. 2b). To monitor the same ALM neurons across days, we saved 6 reference images with 10 µm interval around the most superficial imaging plane for all imaging sessions and identified the most similar imaging plane based on visual inspection across sessions.

Multiple FOVs were imaged across multiple days in each task context. The same set of FOVs were imaged across multiple task contexts. Across all experiments, the total duration from the first imaging session to the last imaging session was 26-233 days (Extended Data Fig. 1g; 95.86 ± 71.95 days, mean ± SD across mice).

### Behavior data analysis

Performance was computed as the fraction of correct choices, excluding early lick trials and no lick trials. Mice whose performance never exceeded 70% after 35-40 days of training were considered unsuccessful in task learning (Extended Data Fig. 1h-i). Chance performance was 50%. Behavioral effects of photoinhibition were quantified by comparing the performance under photoinhibition with control trials using paired two-tailed t-test (Fig. 1i). To quantify the speed of task learning in a given task context (Fig. 1f, Extended Data Figs. 1c, 6g, and 10d), we calculated the number of trials to reach performance criteria of >75% correct and <50% early lick for 100 trials. We excluded the trials in the head-fixation training protocol from the initial task learning for a fair comparison.

### Video data analysis

We used DeepLabCut ^100^ to track manually defined body parts. Separate models were used to track tongue and jaw movements (Extended Data Fig. 1a-b). The development dataset for model training and validation contained manually labeled videos from multiple mice and multiple sessions (correct trials only). For tongue network model, 6 markers were manually labelled in 500 video frames. For jaw network model, 5 markers were manually labeled in 300 video frames. The frames for labeling were automatically and uniformly selected by the program at different timepoints within trials. The labeled frames of the training dataset were split randomly into a training dataset (95%) and a test dataset (5%). Training was performed using the default settings of DeepLabCut. All models were trained up to 500,000 iterations with a batch size of one. The trained models tracked the body features in the test data with an average tracking error of less than 2.5 pixels ^99^.

To analyze tongue and jaw movements during the response epoch, we defined single lick events based on continuous presence of the tongue volume in each frame ^67^. Tongue volume was determined from the internal area of the 4 tongue markers (Extended Data Fig. 1a, left), which were located at the corners of tongue. Lick events were separately grouped based on the lick duration for further time-bin-matched correlation analysis. X and y pixel positions of the tongue tip trajectories were calculated by averaging the frontal tongue markers in each frame. X and y pixel positions of the jaw tip trajectories were calculated by averaging the 3 frontal jaw markers in each frame. For each lick event, we obtained 4 time-series (x position, y position, x velocity, and y velocity) for the tongue (or jaw) tip trajectories (Extended Data Fig. 1a-b, middle). To calculate the similarity between the tongue (or jaw) tip trajectories across lick events (within lick left or lick right), we computed Pearson correlation on the time series for all pairwise lick events within and across sessions. We then calculated the average correlation for the 4 parameters (x position, y position, x velocity, and y velocity) and compared them within session and across sessions (Extended Data Fig. 1a-b, right).

To examine jaw movements during the delay epoch across task contexts, we calculated the x and y displacement jaw tip position by subtracting the average jaw position in a baseline period (1.57 s) before the sample epoch (Extended Data Fig. 6f).

### Preprocessing of two-photon imaging data

Imaging data were preprocessed using Suite2p package ^101^ to perform motion correction and extract raw fluorescence signals (*F*) from automatically identified regions of interest (ROIs). ROIs with >1 skewness were used for further analyses. Neuropil corrected trace was estimated as *F*_neuropil_corrected_(t) = *F*(t) – 0.7 x *F*_neuropil_(t). To visualize activity (Fig. 1d top and Extended Data Fig. 2j left), Δ*F*/*F*_0_ (type 1) was separately calculated in each trial as (*F*−*F*_0_)/*F*_0_, where *F*_0_ is the baseline fluorescence signal averaged over a 1.57 s period immediately before the start of each trial. For all other analyses, we calculated deconvolved activity to avoid the spillover influence of slow-decaying calcium dynamics across task epochs (Extended Data Fig. 2j). To calculate deconvolved activity, *F*_neuropil_corrected_ from all trials were concatenated and Δ*F*/*F*_0_ (type 2) was calculated as (*F*−*F*_0_)/*F*_0_, where *F*_0_ is a running baseline calculated as the median fluorescence within a sliding window of 60 s. Subsequently, Δ*F*/*F*_0_ (type 2) was deconvolved using the OASIS algorithm ^72^ (Extended Data Fig. 2j) after estimating the time constant by auto-regressive model with order p=1. Deconvolved activities were used for all the analyses in this study, except in Fig. 2d (top) and Extended Data Fig. 2j (left) where Δ*F*/*F*_0_ (type 1) traces were shown. Type 1 and type 2 Δ*F*/*F*_0_ only differed in their *F*_0_ calculation.

To track the activity of the same neurons across days, spatial footprints of individual ROIs from the same FOVs were aligned across different imaging days using the CellReg pipeline ^71^. This probabilistic algorithm computes the distributions of centroid distance and spatial correlation between neuronal pairs of the nearest neighbor and all other neighbors within a 10 μm distance (Extended Data Fig. 2g-h). Based on the bimodality between distributions (nearest neighbors vs. other neighbors), CellReg algorithm calculates the estimated false positive and false negative probabilities. By minimizing both estimated error rates for each pair of ROIs, this probabilistic algorithm identifies co-registered neurons and quantifies registration scores for these co-registered neurons (Extended Data Fig. 2i). If the mean squared errors of both centroid distance and spatial correlation model are above 0.1 (a pre-determined hyperparameter), CellReg algorithm generates an error and the FOV is considered as a failure to find co-registered neurons across days. One mouse was removed from all subsequent neuronal data analyses due to failures to find matched neurons across days from all imaging sessions, primarily due to poor imaging window quality. Among co-registered neurons, only neurons with reliable responses in at least one imaging session (i.e., Pearson correlation between trial-averaged and trial-type-concatenated Δ*F*/*F*_0_ (type 1) PSTHs calculated using the first versus second halves of the trials >0.5) were used for further analyses.

In the experiment where we imaged the same FOV across multiple sessions in the same task context, we define the sessions as expert-early and expert-late sessions (Fig. 2). In cases where we imaged the same FOV twice over time, the 2 sessions were defined as expert-early and expert-late sessions accordingly. In cases where we imaged more than 2 sessions from the same FOV over time, the expert-early and expert-late sessions were defined for pairs of sessions.

Specifically, for single neuron analyses (e.g. Figs. 2e and 2k), we only compared the first and second imaging sessions to avoid inclusion of duplicate data points from the same session. These two sessions are defined as expert-early and expert-late sessions, respectively. For population level activity projection and decoding analyses (Fig. 2i-j), we included all the possible pairwise comparisons. For each pair, the two sessions used are defined as expert-early and expert-late sessions, respectively.

### Two-photon imaging data analysis

Neurons were tested for significant trial-type selectivity during the sample, delay, and response epochs, using deconvolved activities from different trial types (non-paired two-tailed t-test, P<0.001; correct trials only). We used the early sample epoch (first 0.83 s, 5 imaging frames), late delay epoch (last 0.67 s, 4 frames), and early response epoch (first 1.33 s, 8 frames) as the respective time windows for the statistical comparisons and all the following analyses (Extended Data Fig. 4a-c). To examine the stability of single neuron selectivity index, we first identified significantly selective neurons in each task epoch. We then determined each neuron’s preferred trial type (“lick left” versus “lick right”) using the earlier imaging session in task context 1. Next, selectivity index was calculated as the difference in activity between trial types divided by their sum (anterior versus posterior pole position for sample epoch selectivity; lick left versus lick right for delay and response epoch selectivity; correct trials only). To define preferred trial types in earlier sessions, a portion of the trials were used for statistical tests to determine significant selectivity and the preferred trial type, then independent trials were used to calculate selectivity index within the same session. We then calculated selectivity for the defined neurons in later sessions or across different task contexts.

For error trial analysis (Extended Data Fig. 2k-l), only the imaging sessions with more than 10 error trials for each trial type were analyzed. Selectivity was calculated as the difference in trial-averaged activity (deconvolved calcium activity) between instructed lick right and lick left trials, using correct and error trials separately. Selectivity was calculated during the early sample epoch, late delay epoch, and response epoch.

To analyze the encoding of trial types in ALM population activity, we built linear decoders that were weighted sums of ALM neuron activities to best differentiate trial types. We examined the encoding of four kinds of trial types: 1) anterior versus posterior pole position trials for stimulus encoding during the sample epoch in the tactile-instructed lick task; 2) low tone (2 kHz) versus high tone (10 kHz) for stimulus encoding during the sample epoch in the auditory-instructed lick task; 3) lick left versus lick right for lick direction encoding during the delay epoch; 4) lick left versus lick right for lick direction encoding during the response epoch.

To build the linear decoder for a population of *n* ALM neurons, we found a *n* x 1 vector coding direction (**CD**) in the *n* dimensional activity space that maximally separates response vectors in different trial types during defined task epochs, i.e., **CD_Sample_** for stimulus encoding during the sample epoch, **CD_Delay_** for lick direction encoding during the delay epoch, and **CD_Response_** for lick direction encoding during the response epoch. To estimate the **CD**’s, we first computed **CD***_t_* at different time points as:

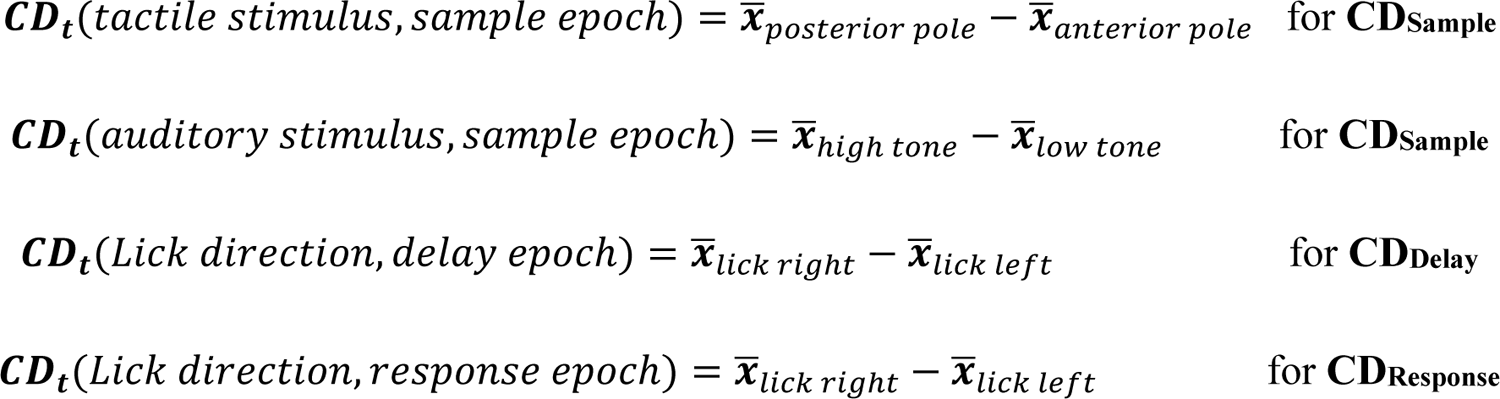

where *x*^-^’s are *n* × 1 trial-averaged response vectors that described the population response for each trial type at each time point, *t*, during the defined task epochs. Next, we averaged the **CD***_t_*’s within the defined task epoch to separately estimate the **CD_Sample_**, **CD_Delay_**, and **CD_Response_**. **CD_Sample_**, **CD_Delay_**, and **CD_Response_** were computed using 50% of trials and the remaining trials from the same session or from different sessions were used for activity projections and decoding (Fig. 2g; correct trials only).

To project the ALM population activity along the **CD_Sample_**, **CD_Delay_**, and **CD_Response_**, we computed the deconvolved activity for individual neurons and assembled their single-trial activity at each time point into population response vectors, ***x****’s* (*n* x 1 vectors for *n* neurons). The activity projection in Figs 2-5 and Extended Data Figs. 3-5, 7, 9 were obtained as **CD** ^T^***x***, **CD** ^T^***x***, and **CD_Response_**^T^***x***.

To decode trial types using ALM population activity projected onto the **CD_Sample_**, **CD_Delay_**, and **CD_Response_** (Figs. 2-5 and Extended Data Figs 4, 5, 7, 9), we calculated ALM activity projections (**CD_Sample_**^T^***x***, **CD_Delay_**^T^***x***, and **CD_Response_**^T^***x***) within defined time windows and we computed a decision boundary (DB) to best separate different trial types:

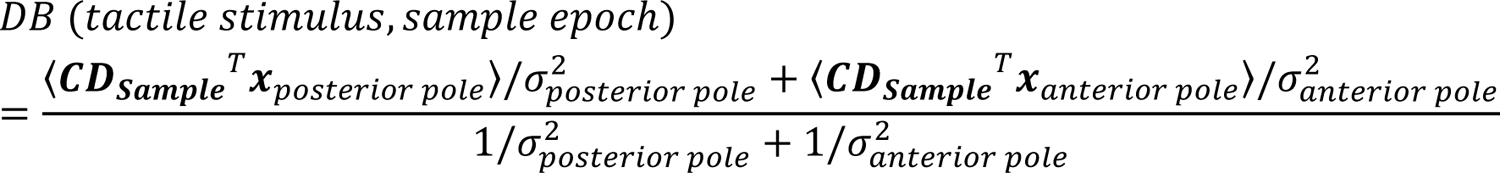

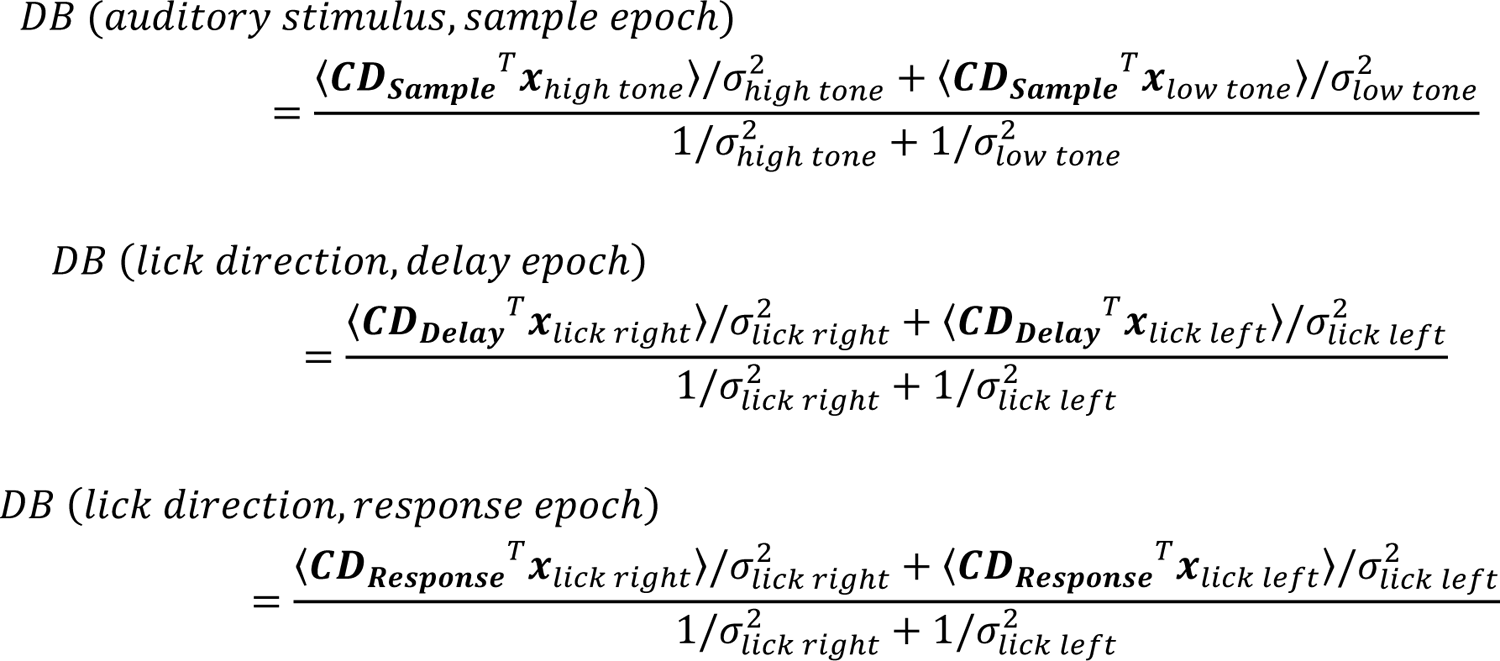

σ^2^ is the variance of the activity projection ***CD***^*T*^*x* within each trial types. DB’s were computed using the same trials used to compute the **CD**’s and independent trials were used to predict trial types. To examine decoding performance across task contexts, we restricted the analysis to decoders with accuracy of > 0.7 within the session it was trained in (cross-validated performance). This is because if a decoder exhibited low decoding performance to begin with, its decoding performance will be generally low in other sessions due to poor training of the decoder.

To analyze activity changes along other dimensions of activity space across task contexts, we defined a ‘uniform shift (**US**) axis’ ^11^ using trial-type-averaged activity:

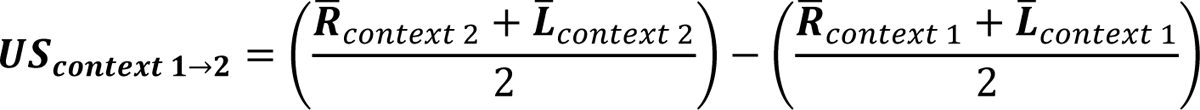

where *R*^-^ and *L*^-^ are *n* × 1 response vectors that described the trial-averaged population response for lick left and lick right trials at the end of the delay epoch. We separately calculated **US**’s for each task context change, i.e., **US_1_**→**_2_** for task context 1→2, **US_2_**→**_1’_** for task context 2→1’,

**US_1’_**→**_2’_** for task context 1’→2’ (Extended Data Fig. 8b). For activity projections (Extended Data Fig. 8c), the **US**’s are further orthogonalized to the **CD**’s using the Gram-Schmidt process to capture activity changes along dimensions of activity space that were not selective for lick direction (‘movement-irrelevant subspace’). We computed the **US**’s using 50% of the trials and the remaining 50% of the trials were used for activity projections (Extended Data Fig. 8c). The dot products in Extended Data Fig. 8d were calculated without any orthogonalization.

### Modeling

The instructed directional licking task with a delay epoch was modeled with simulations lasting for two seconds. The first second of the simulation was the sample epoch during which time trial-specific external inputs were provided and the last second was the delay epoch in which the inputs were removed. The coding direction, ***CD***_*Delay*_ was calculated as the difference between network activity on lick left and lick right trials at the end of the delay epoch (*t* = 0), similar to the neural data. The trial type was always defined by instructed lick direction in different task contexts (across contingency reversals).

### Recurrent Neural Networks

Recurrent neural networks (RNNs) consisted of 50 units with dynamics governed by the equations

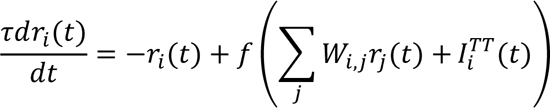

where *r*_*i*_(*t*) is the spike rate of neuron *i*, the synaptic time constant τ was set equal to 200 ms,

*W*_*i,j*_ is the synaptic strength from neuron *j* to neuron *i, T*^*TT*^_i_(*t*) is the trial-type (*TT*) dependent external input to neuron *i*, and *f*(*x*) = tanh (*x*) is the neural activation function.

The connection matrix *W* was randomly initialized from a Gaussian distribution. The network was scaled to have a maximum eigenvalue equal to 0.9. To generate persistent activity, networks must have an eigenvalue greater than or equal to one. Networks initialized with eigenvalues greater than one tended to learn the task with high dimensional persistent activity, inconsistent with ALM dynamics ^19^. Initializing with eigenvalues less than one tended to produce lower dimensional persistent activity.

External input strengths *T*^*TT*^ were drawn from a Gaussian distribution with mean equal to zero and standard deviation of 0.3. Two distinct input vectors were used for anterior *I*^*A*^_i_ and posterior *I*^*P*^_i_ pole position trials.

Behavioral readout *B* was given by the linear projections 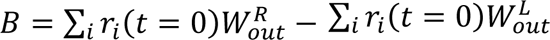, where *t* = 0 is the time at the end of the delay epoch, *W*^*R*^_*Out*_ and *W*_*Out*_^*L*^ are gaussian random readout vectors corresponding to rightward and leftward movements, respectively.

RNNs were trained using backpropagation through time (BPTT). The input (*T*^*TT*^) and readout weights (*W*_*Out*_^*R*^ and *W*_*Out*_^*L*^) were fixed and only the recurrent weights *W*_*i,j*_ internal to the RNN were trained. For each trial type, activity along the correct readout direction was trained to match a linear ramp of activity starting at the beginning of the sample epoch and the incorrect readout direction was trained to have zero activation. For task context 1, presentation of *T*^*A*^ was associated with ramping along *W*_*Out*_^*L*^ and zero activation along *W*_*Out*_^*R*^, presentation of *T*^*P*^ was associated with the opposite behavior. These associations were reversed for task context 2. Networks were trained for 100 iterations.

In the RNNs, the behavior readout relied on many units (dense *W*_*Out*_^*R*^ and *W*_*Out*_^*L*^). Because only 2 units in the AFF networks contributed to behavior output, this difference in readout may affect how these networks learned to produce reversed output. We therefore also tested RNNs in which we fixed the behavior readout to only 2 units like the AFF network (sparse *W*_*Out*_^*R*^ and *W*_*Out*_^*L*^), but all results remained unchanged.

### Amplifying feedforward network

ALM circuitry contains an amplifying feedforward (AFF) circuit motif ^77^. The AFF network is a recurrent circuit in which preparatory activity during the delay epoch flows through a sequence of activity states. Each activity state can be modeled as a layer within a feedforward network. In addition, the late layers in the network are connected to early layers through feedback connections. Here we develop a framework for training AFF networks to generate choice-selective persistent activity.

Before detailing the learning rules used for training AFF networks, we first introduce several features that make AFF networks advantageous for training. Training neural networks require pathways linking input units to output units for computation, and pathways linking outputs to inputs for learning. In the simplest cases, output to input feedback may interfere with the input to output computations. AFF networks, and non-normal networks in general, do not generate reverberating feedback. For this reason, it is possible to construct AFF networks that bidirectionally link inputs to outputs through separate channels that do not interfere with each other.

Amplifying feedforward (also commonly referred to as non-normal) networks are constructed by applying orthonormal transformations to purely feedforward networks. Orthonormal transformations to feedforward networks serve two useful anatomical purposes: i.) they form feedback connections from late layers to early layers and ii.) they form stabilizing excitatory/inhibitory connections to eliminate any reverberation that may result from the newly formed feedback connections. In this model, we use the feedback connections from late layers to early layers to convey performance feedback signals allowing the AFF network to learn via error backpropagation.

We first constructed a purely feedforward network with 4 layers referred to as Input (n; 30 units), hidden layer 1 (h1; 200 units), hidden layer 2 (h2; 5 units) and output (o; 2 units) (Extended Data Fig. 11). Trial-type (*TT*) dependent external inputs, *T*^*TT*^(*t*), were provided only to the input layer.

Feedforward connection matrices (*W*_*i,j*_^*n,h*1^, *W*_*i,j*_^*h*1,*h*2^ and *W*_*i,j*_^*h*2,o^) conveyed these inputs to downstream layers and were initialized from a uniform positive distribution. Next, we added feedback connections from o to h2 (*W*_*i,j*_^o,*h*2^) and from h2 to h1 (*W*_*i,j*_^*h*2,*h*1^) to provide performance feedback for training the feedforward connections. Feedback connections were matched to feedforward connections so that *W*_*i,j*_^o,*h*2^ = *W*_*i,j*_^*h*2,o^. These feedback connections provide scaffolding to precisely implement error backpropagation to train feedforward connections. However, the presence of feedback connections in the circuit will introduce feedback to the network that will interfere with its feedforward computations.

To cancel out the reverberations caused by this feedback we incorporated additional stabilization hidden layers s1 (200 units) and s2 (5 units) (Extended Data Fig. 11). Each hidden unit in layer h1 is matched with a stabilizing neuron in the stabilization layer s1 which receives the same feedback connections as its paired excitatory neuron and projects inhibitory connections of the same strength as its excitatory partner. Similarly, each neuron in h2 has a corresponding unit in s2. Mathematically this relationship is written as

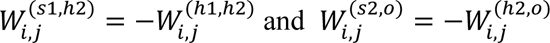

And

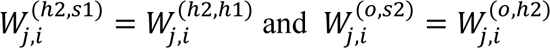

Because of the precisely balanced excitation and inhibition, this recurrent network is non-normal; all eigenvalues are equal to zero. This non-normal network has two independent pathways, one linking the input layer to the output layer, useful for computation; and the other linking the output layer to the input layer, useful for learning.

The network is trained using error backpropagation; an error signal is computed and then sent back into each unit in the output layer. This error signal is conveyed to the early layers by the feedback connections. The stabilizing network ensures that this error signal does not reverberate. The backpropagated signal in neuron *i* in the hidden layers *h*1 and *h*2 are thus given by the equations

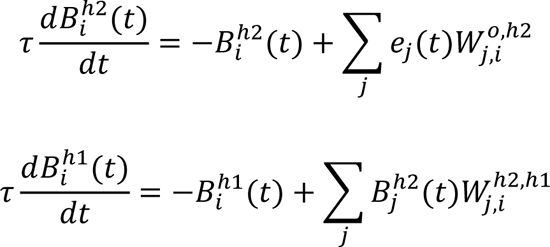

As in error backpropagation, feedforward weights (i.e. *W*^(*h*1,*h*2)^) are updated by taking the product of the forward pass activity and the backward pass activity. For example, connections from neuron *i* in layer *h1* onto neuron *j* in layer h2 are updated according to the rule

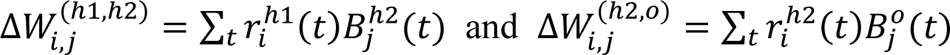

This rule is applied to all feedforward connections (i.e. *n* → *h*1, *h*1 → *h*2, and *h*2 → o). Changing the feedforward weights will necessarily disrupt the precise balance in the network. To maintain stability, the stabilizing weights must be updated to precisely cancel the changes to the feedforward weights

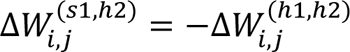

Compensatory weight changes based on this equation are applied to all connections in the stabilization layers (i.e. *S*1 → *h*2 and *S*2 → o).

The AFF network was trained to form the same associations as the RNN. Unlike the RNN, the AFF utilized a linear neuronal activation (*f*(*x*) = *x*) so that dynamics are governed by the equation

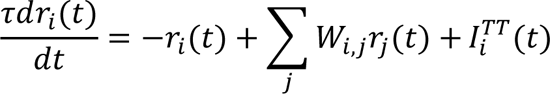

Additionally, because the AFF naturally generates ramping signals^77^, the output units were not trained to match a ramping signal at all time points, but rather trained to be activated at a specific level at the end of the delay. For example, the target for the lick right output unit (*T*_*R*_) on posterior trials was *T*_*R*_(*t* = 0) = 6 and *T*_*R*_(*t* = 0) = 0 on anterior trials.

### Analysis of neural dynamics within RNN and AFF networks

For each network, we calculated the selectivity of each unit as the activity difference between the lick right and lick left trials in each task context. We calculated eigenvectors of the network selectivity matrix using singular value decomposition (SVD). The data for the SVD was an *n* × *t* matrix containing the selectivity of *n* units over *t* time bins (selectivity from task contexts 1 and 2 were concatenated). 3 vectors usually captured most of the network activity variance across both task contexts (Extended Data Fig. 11f). We then rotated the 3 eigenvectors so that the first vector was aligned to the dimension that maximized the difference in network selectivity matrix between task contexts 1 and 2. Network activity projected on the first vector was correlated with the network input across task contexts, thus referred to as the stimulus mode (Extended Data Fig. 12a-b). Network activity projected on the second vector was correlated with the network output across task contexts and exhibited ramping activity during the delay epoch, thus referred to as the output mode (Extended Data Fig. 12a-b).

To examine the **CD_Delay_** reorganization across task contexts as a function of stimulus mode strength (Extended Data Fig. 12c), we summed the network activity projected on the stimulus mode across time. This activity strength was normalized to the mean activity of each network to enable comparisons across different networks.

### Statistics

The sample sizes were similar to sample sizes used in the field: for behavior and two-photon calcium imaging, three mice or more per condition. No statistical methods were used to determine sample size. All key results were replicated in multiple mice. Mice were allocated into experimental groups according to their strain or by experimenter. Unless stated otherwise, the investigators were not blinded to mouse group allocation during experiments and outcome assessment. Trial types were randomly determined by a computer program. Statistical comparisons using *t*-tests and other statistical tests are described above. We used Pearson’s correlation for the linear regression. Error bars indicate mean ± s.e.m. unless noted otherwise.

## Code availability

Code used for data analysis is available at https://github.com/NuoLiLabBCM/KimEtAl2024.

## Data availability

Raw and processed data are available from the corresponding author upon request.

